# Using Machine Learning to identify microRNA biomarkers for predisposition to Huntington’s Disease

**DOI:** 10.1101/2022.08.16.504104

**Authors:** K Patel, C Sheridan, DP Shanley

## Abstract

**Background:** Huntington’s disease (HD) is an autosomal dominant disease which is triggered by a large expansion of CAG nucleotides in the *HTT* gene. While the CAG expansion linearly correlates with the age of disease onset in HD, twin-studies and cohorts of Juvenile Onset HD (JOHD) patients have shown other factors influence the progression of HD. Thus, it would be of interest to identify molecular biomarkers which indicate predisposition to the development of HD, and as microRNAs (miRNAs) circulate in bio-fluids they would be particularly useful biomarkers. We explored a large HD miRNA-mRNA expression dataset (GSE65776) to establish appropriate questions that could be addressed using Machine Learning (ML). We sought sets of features (mRNAs or miRNAs) to predict HD or WT samples from aged or young mouse cortex samples, and we asked if a set of features could predict predisposition to HD or WT genotypes by training models on aged samples and testing the models on young samples. Several models were created using ADAboost, ExtraTrees, GaussianNB and Random Forest, and the best performing models were further analysed using AUC curves and PCA plots. Finally, genes used to train our miRNA-based predisposition model were mined from HD patient bio-fluid samples.

**Results:** Our testing accuracies were between 66-100% and AUC scores were between 31-100%. We generated several excellent models with testing accuracies >80% and AUC scores >90%. We also identified homologues of *mmu-miR-154-5p*, *mmu-miR-181a-5p*, *mmu-miR-212-3p, mmu-miR-378b, mmu-miR-382-5p* and *mmu-miR-770-5p* from our miRNA-based predisposition model to be circulating in HD patient blood samples at p.values of <0.05.

**Conclusions:** We generated several age-based models which could differentiate between HD and WT samples, including an aged mRNA-based model with a 100% AUC score, an aged miRNA-based model with a 92% AUC score and an aged miRNA-based model with a 96% AUC score. We also identified several miRNAs used to train our miRNA-based predisposition model which were detectable in HD patient blood samples, which suggests they could be potential candidates for use as non-invasive biomarkers for HD research.

## Introduction

Huntington’s Disease (HD) is a rare autosomal dominant disease, characterised by the progressive destruction of cortical and striatal neurons. HD patients have decreased motor skills, memory, and overall ability to maintain themselves without aid ^1^. HD affects 9.3-13.7 individuals per 100,000 in western populations ^2,3^. Another form of the neurodegenerative disorder is Juvenile Onset Huntington’s Disease (JOHD) which affects 1-9.6% of reported HD cases ^4^. CAG nucleotide repeats in the Huntingtin gene (*HTT*) leads to the translation of mutated HTT (mHTT) proteins with large a polyglutamine repeat chain. Genetic screening of the CAG repeats is performed regularly as the length of the CAG repeats inversely correlates with adult-onset HD. Having <27 CAG repeats is indicative of no penetrance of HD, 27-35 CAG repeats indicate an intermediate chance of penetrance, >36 CAG repeats will lead to full penetrance of HD, and patients with >60 CAG repeats develop JOHD ^2,5,6^. HD patients develop disease phenotypes during middle-aged life and JOHD patients will show symptoms prior to age 25 ^1^. Understanding the underlying mechanisms of the HTT protein can help to investigate how the neurotoxicity from HD is triggered. HTT is found in the cytoplasm of striatal and cortical neurons ^7^. Many HEAT binding motifs are found on HTT implying it has roles as a scaffolding protein and is involved in many protein interactions ^8^. However, a large CAG nucleotide expansion in the mHTT protein halts its ability to exit the nucleus and thus mHTT is incapable of interacting with other scaffolding proteins: B-tubulins, microtubulins, dynein, dynactin ^9–11^. Gene expression changes indicate that HTT has roles as a transcriptomic regulator. A well characterised example is normal HTT proteins bind and sequester REST (repressor element-1 transcription factor) a negative regulator of BDNF (brain-derived neurotrophic factor) which is a neuron survival factor. mHTT cannot sequester REST, leading to decreased BDNF levels ^12,13^.

Currently no therapies for HD are available, and though age is a key risk factor further mechanistic knowledge is needed ^14,15^. This was highlighted by twin studies where patients with the same genotype did not have uniform experiences HD ^15,16^. Furthermore, in several JOHD cohorts non-linear correlations were identified between the length of CAG expansion which and age of onset of the disease ^17–19^. Better understanding of genetic and epigenetic factors could lead to novel insights for HD research. Transcriptomic studies have shown mHTT changes expression profiles of neuronal of mRNAs and non-coding RNAs such as microRNAs (miRNAs) ^20,21^. miRNAs are small (18-22 nucleotides) single stranded RNAs which negatively regulate specific mRNAs via complementary binding ^22^. miRNAs contain a 6-8 nucleotides long tract called the seed site on their 5’ end which complementary binds to target sites on the 3’-UTR of the mRNA ^23^. Biogenesis of miRNAs is a tightly regulated process which begins with transcription via RNA polymerase II ^24^.

This produces a pri-miRNA (70-100 nucleotide long double stranded RNA hairpin) which will undergo further modifications by nuclear proteins DROSHA and DGCR8 ^25^. A pre-miRNA hairpin will be formed, and it will be detected as cargo by EXP5 for transportation to the cytoplasm ^26^. Cytoplasmic proteins DICER and TRBP perform further biogenesis and recruit the RISC complex which will attach itself to a mature miRNA strands which will act as the guide for the RISC complex ^27,28^. The RISC complex comprises of sub-units such as AGO2 and a GW182 protein, and in mammals the GW182 protein is commonly responsible to mRNA silencing by orchestrating de-capping, de-adenylation and 5’-3’ decay events (Figure 1) ^28–32^. Both the seed site of the miRNA and the binding sites found on the 3’-UTR of target mRNAs are highly evolutionarily conserved ^33^. A major interest for neurodegenerative research in miRNAs is their ability to shuttle in bio-fluids as they could be used as non-invasive biomarkers which can be measured in blood plasma or cerebrospinal fluid (CSF) to assess the health of brains ^34,35^.

**Figure 1.**
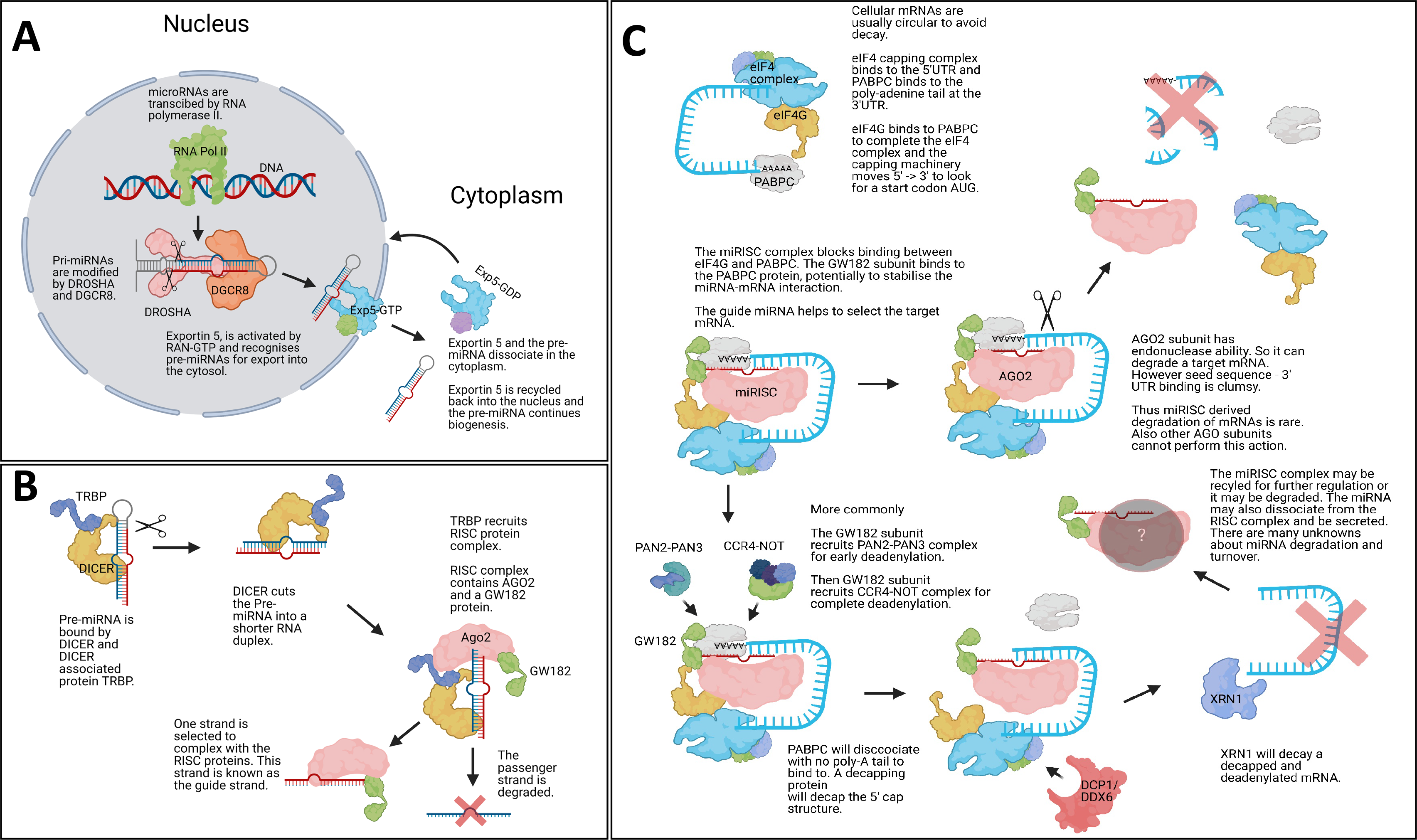
miRNA biogenesis. The Illustrations show canonical mammalian miRNA biogenesis steps. (**A)** Shows the initial transcription of the mature miRNA via RNA polymerase II and subsequent processing by DROSHA and DGCR8, which creates a pre-miRNA stem-loop structure which is exported into the cytoplasm by Exportin5. (**B)** Within the cytoplasm DICER and TRBP cut the loop of the pre-miRNA and the RISC complex is associated with the mature guide miRNA strand. The passenger strand is degraded. (**C)** The newly form miRISC complex will interfere with the eukaryotic initiation machinery via the GW182 protein binding to the PABPC protein. GW182 protein will then recruit deadenylation complexes, and this is followed by a decapping protein removing the CAP protein complex from the target mRNA. The naked mRNA structure is decayed by XRN1 and the miRISC complex may be recycled. Some mammalian miRNA-based mRNA degradation is based on exonuclease activity by AGO2. Image was made with BioRender.

In this paper we used machine learning (ML) to identify biomarkers for HD from young and aged samples and we also used ML to predict predisposition of young samples to HD (Figure 2). A valuable dataset to explore potential biomarkers was stored in GSE65776. This dataset contained 168 RNAseq and miRNAseq samples from male and female mouse cortexes, and the mice were sacrificed at 2, 6 and 10 months old (M). Most samples had knock-in mutations which would lead to JOHD in humans, and these were: 80, 92, 111, 140 and 175 CAG repeats in the *HTT* gene. The authors of the dataset reported that the 80 – 111 CAG repeat mice had molecular changes associated with HD but no/ mild visible phenotype and the 140 – 175 CAG repeat mice had phenotypes which reflected human HD. This dataset also included WT samples and control samples which had 20 CAG repeat knock-ins ^20,21^. The initial data generation and analysis paper showed striatal neuron samples had greater gene expression changes than cortical neuron samples, however we were more interested in cortical samples as cortical loss has been recorded as an early pathological event of HD and this would be useful for our early predisposition queries ^36^. Initial data exploration with differential expression (DE) analysis and the *TimiRGeN R* package identified a batch effect in the data, high homology, and some gender differences ^37^. Based on this, the data was corrected (batch effect removed, gender differences reduced) and re-labelled so that HD and WT samples could be differentiated using ML. While miRNAs and mRNAs were kept separated, three lines of enquiry were established: identification of post-symptomatic markers in aged (10M) samples, identification of pre-symptomatic markers in young (2M) samples, and identification of predisposition markers by training on aged samples and testing on young samples.

**Figure 2.**
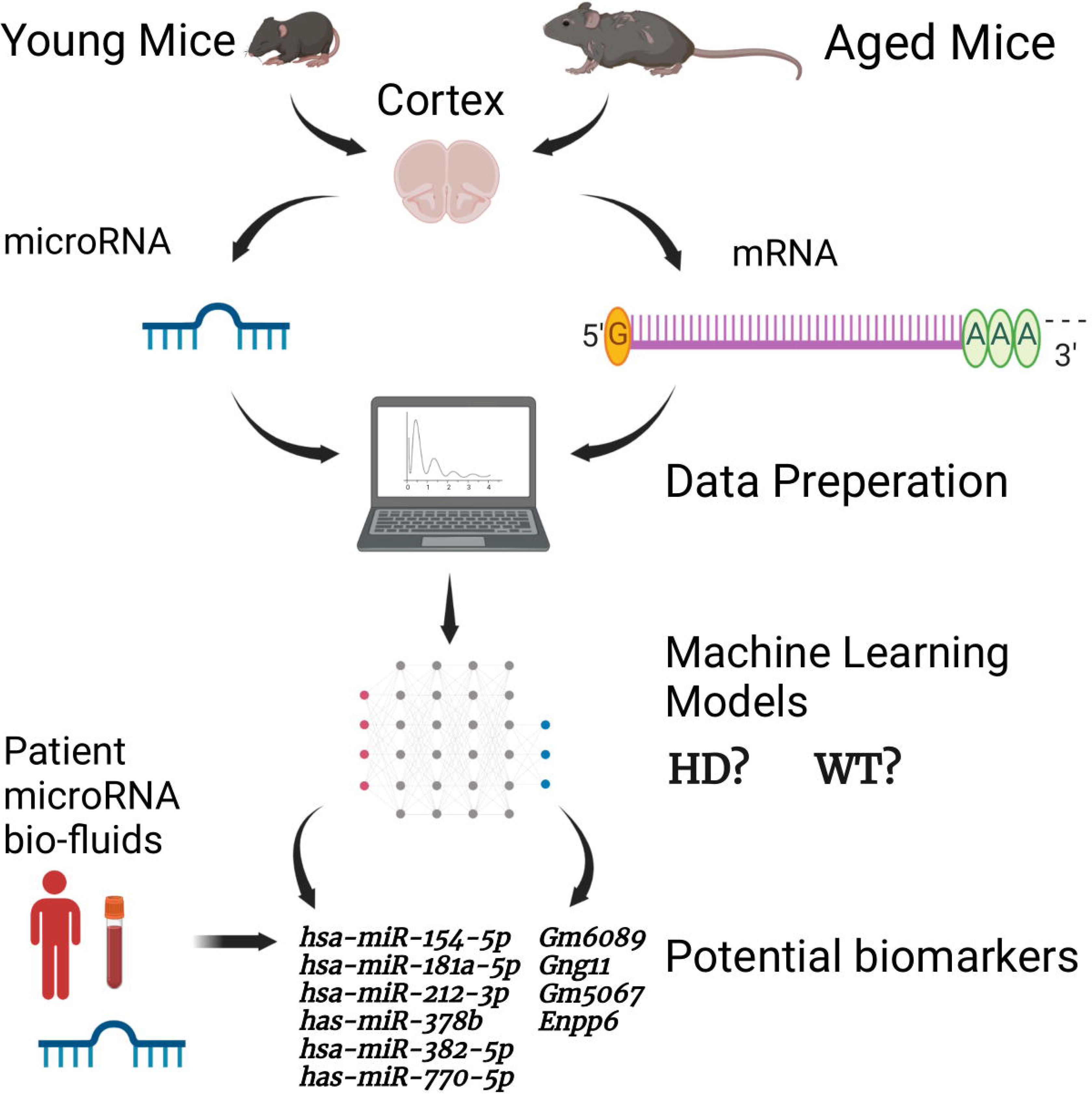
Simplified diagram of the HD ML project presented. The illustration shows the major steps of the project, including how the datasets used in this project were created, the separation of miRNAs and mRNAs, the data processing and ML steps. The image also shows the miRNA and mRNA biomarkers found through the predisposition models. Image was made with BioRender.

## Materials and Methods

### Data download and raw data pre-processing

Fastq files were downloaded from GSE65776 (GSE65770 - mRNA and GSE657679 - miRNA) using *SRA-toolkit* ^38^. Quality of the fastq files were checked using *FASTQC* ^39^. For miRNA pre-processing, Mus_musculus.GRCm38.cdna.all.fa was used to create index files with *Bowtie* and mature miRNA count calling was performed with *miRDeep2* ^40,41^. mRNA samples were aligned into bam files using *Salmon* and bam index files were created with *Samtools*. The Mus_musculus.GRCm38.cdna.all.fa transcriptome was used for mRNA alignment ^42^. *Tximport* was then used to aggregate mRNA data to gene count levels ^43^.

### Initial data exploration

The mRNA and miRNA data consisted of 168 samples each. Seven genotypes were present: WT, 20 (control), 80, 92, 111, 140 and 175 CAG repeats. The data consisted of three age groups: 2M, 6M, 10M. Male and female samples were evenly distributed for most genotypes. Outliers were detected using PCA and removed, and lowly expressed genes were removed if their value was <10 in at least 1/3^rd^ (56) of the total number of samples leaving 332 miRNAs and 13715 mRNAs. With *DESeq2* gender and age matched DE contrasts were made between 20 CAG/WT, 80 CAG/WT, 92 CAG/WT, 111 CAG/WT, 140 CAG/WT, 175 CAG/WT ^44^. Genes with a Benjimini-Hochberg FDR of <0.05 were kept. Significantly expressed genes from the gender based DE analysis was taken forward for miRNA-mRNA integrated analysis with the *TimiRGeN R* package for pathway enrichment analysis ^37^.

### Batch removal and removing variance from gender differences

A batch effect in the 6M for the miRNAs and mRNAs was suspected after DE analysis so the 6M was removed (Supplementary Figure 1). Variance from gender differences were reduced with *combat* ^45^. One 10M sample was removed by pca analysis, as it was not within 6* standard deviations from the mean.

### Data preparation prior to ML

We found this data to be highly homogenous based on the lack of differentially expressed genes (DEGs) found when contrasting different CAG lengths with the WT samples, so we increased our sample power by labelling all Q80, Q92, Q111, Q140 and Q175 samples as “HD” samples, and the WT and Q20 samples were labelled as “WT” samples. Male and female samples from the same mutation categories were also labelled together after correcting for gender-based variance. After the correction steps, samples were re-processed to filter for low varying genes. For the miRNA and the mRNA data, genes were removed if their expression was <10 in at least half (56) of the samples. Leaving 519 and 16432 genes for the miRNA and mRNA data respectively. The mRNA data was further filtered to only contain genes which showed a log2fc >0.2 or <−0.2 (after contrasting HD and WT samples with DE), leaving a total of 532 mRNAs. Normalised gene expression levels were extracted from *DESeq2* normalised counts.

### Identified questions to investigate with ML

In line with data centric AI, we identified appropriate questions based on our data. We were interested in six questions: a) which miRNAs are the best set of features to detect HD or WT samples after the onset of symptoms (10M), b) which miRNAs are the best set of features to detect HD or WT samples prior to the onset of the symptoms (2M), c) using miRNAs can we use the aged samples (10M) to identify the best set of features to detect predisposition to HD or WT in young samples (2M), d) which mRNAs are the best set of features to detect HD or WT samples after the onset of symptoms, e) which mRNAs are the best set of features to detect HD or WT samples prior to the onset of the symptoms, f) using mRNAs can we use the aged samples to identify the best set of features to detect predisposition to HD or WT in young samples.

### Data processing for ML and model performances

ML was performed with scikit-learn 2.2.4 ^46^. miRNAs and mRNAs were analysed separately. 10M and 2M data were separated and split into training and testing sets at a 0.8:0.2 ratio, without shuffling. SMOTE was used to create random synthetic WT samples for each training step (but not for testing), and with this we synthesized an additional 24 WT samples for the young dataset and 23 WT samples for the aged dataset (because an outlier was removed from the 10M data) ^47^. Min_max scaling was performed on training and test data. 5-fold cross validations were performed during all model training steps. Recursive feature elimination (RFE) was performed using a linear SVM algorithm and a 5-fold cross-validation at an attempt to remove low variance genes from the training data. Eight popular classifiers were trained and tested, using the default settings for each algorithm, and these were: LinearSVM, PolynomialSVM, Guassian Process, Extra Trees (ET), Random Forest (RF), Neural Network, Adaboost (ADA), Gaussian Naïve Bayes (GNB). We found RFE led to overfitting. A k-best feature selection approach was instead utilised to identify if similar or better predictions could be generated with fewer genes. The four best performing classifiers (ADA, ET, GNB RF) for the miRNA data from RFE were taken forward to determine the optimal features in a robust k-best selection approach. The 1-100 best features found by the k-best method (highest F-score) were sequentially used to train models using each of the four classifiers. This process was repeated 100 times, resulting in 100 × 100 × 4 (40,000) trained models per ML question. To limit the randomness of some classifiers, the best three training accuracies from each classifier were taken forward to create hyperparameter tuned models (Supplementary Table 1). For ADA number of estimators and learning rate were parametrised, for ET number of estimators and minimum splits were parameterised, for GNB the use of an automated variance smoother was contrasted against no variance smoother and if the results were the same – the scores from no variance smoother selected, and for RF number of estimators was parameterised. Training accuracies, testing accuracies, precision scores, recall scores, f1 scores and confusion matrices were calculated for each model (Supplementary Table 2). ROC_AUC and PCA plots were made for the best performing model from each of the six questions (Figure 3). miRNA-based feature selection and model building were reperformed using only Q140 and Q175 as the HD samples. Reperformance data comprised of 16 WT and 16 HD samples for the 2M data and 16 WT and 15 HD samples for the 10M data. K-best feature selection and hyperparameter tuning was carried out as above, and SMOTE, min_max scaling and a 5-fold cross validation were applied. Since the classes were more balanced SMOTE had a limited effect as it only synthesised 1 HD sample for the 10M data. While TP = true positives, TN = true negatives, FP = false positives and FN = false negatives, the equations of importance are displayed below.

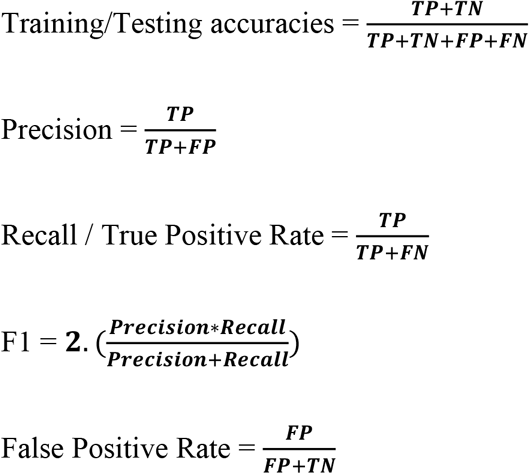

**Figure 3.**
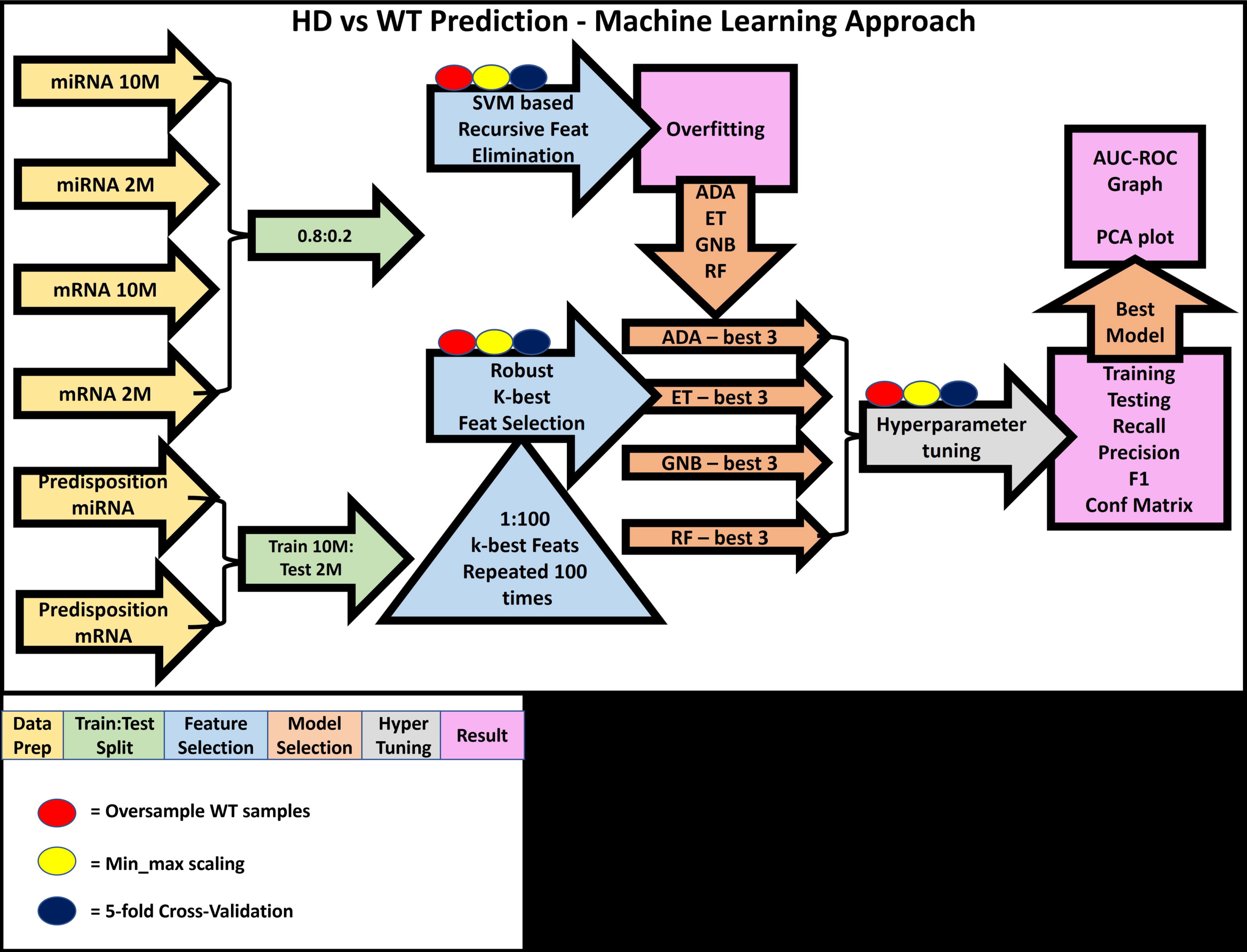
Machine learning approach. This is an illustration our ML approach, including how the data was prepared (yellow arrows) for each of the six questions and how the data was split (green arrows). For the age matched classification questions, a simple 0.8:0.2 training to testing split was used, and for the more ambitious predisposition questions the 10M data was trained on and the 2M data was tested on. Following this, the RFE and k-best based feature selection (blue arrows and blue triangle) were performed. RFE was performed first and from its results (pink boxes) we were able to inform our k-best strategy by using classifiers which performed best on the miRNA-based ML questions (orange arrows, Supplementary Table 1). The three best performing models (orange arrows) from each classifier were hyperparameter tuned (grey arrow), and performance scores were measured (Supplementary Table 2). For each question, the best performing model had AUC-ROC curves and PCA plots made for them. For all model training steps, oversampling of WT samples (red circles), min_max scaling (yellow circles) and a 5-fold cross-validation strategy (dark blue circles) was used. Oversampling and 5-fold cross validation were only performed with training samples to limit leakage, and scaling was performed on training and test data.

### Systematic check in bio-fluid datasets

Processed samples were downloaded from GSE167630, GSE108395 and GSE108396 ^48,49^. Each were adult-onset HD vs adult control datasets which were either created using human miRNA specific microarrays or microRNA-seq. GSE167630 measured miRNA expression in blood samples, GSE108395 and GSE108396 were part of the same study (GSE108398) and respectively measured miRNAs in blood plasma and CSF. Datasets were checked for normalisation and visual outliers, and then analysed using standard methodology with either *Limma* or *DESeq2* ^44,50^. Using MirBase and TargetScans we found and 35 of the 60 mouse miRNAs used for training the best performing predisposition model had known human homologues ^51,52^. The homologues identified in the miRNA-based predisposition model were mined and then researched for relevant literature.

## Results

### Initial data exploration found high homogeneity between WT and HD samples

Using DE on age and gender matched miRNA samples found between 0-6 DEGs from the 2M samples, between 0-17 DEGs from the 10M samples and surprisingly at least 230 DEGs from the 6M, including when contrasting the positive control 20 CAG samples with the WT samples. A similar trend was seen with the mRNAs as we found between 0-34 DEGs for the 2M samples, between 0-814 DEGs for the 10M samples and over 2000 DEGs in many of the 6M samples. The odd pattern in DE genes (Supplementary Table 3) made us suspect a batch effect in the 6M data, so it was removed from further analysis. The homogeneity of the data and the batch effect can be seen in PCA plots (Supplementary Figure 1). DE also suggested that there may be some differences in how HD develops in different genders, and to further investigate for mechanistic differences between male and female samples we used the *TimiRGeN R* package. DEGs from: male 140 CAG 10M /male WT 10M samples were enriched in Triacyglyceride Synthesis, Sphingolipid Metabolism (integrated pathway) and Sphingolipid Metabolism (general overview), female 175 CAG 2M / female WT 2M samples were enriched in IL-9 Signaling Pathway and female 175 CAG 10M / female WT 10M were enriched in Myometrial Relaxation and Contraction Pathways, Calcium Regulation in the Cardiac Cell, Omega-9 FA synthesis, Glycosis and Gluconeogenesis and Cholestoral metabolism. This informed us that it would be best to reduce variance between the genders. The major conclusion from the initial data exploration was that this data had very low numbers of DEGs (after removing the suspected batch effect) which indicated high homogeneity between WT and HD samples, and so we devised an alternative approach to identify miRNA biomarkers from this data set. Using the clinical cut-offs of <27 CAG repeats leading to no disease and >36 CAG repeats leading to HD, we opted to re-label all samples with CAG repeats of 80 or more as “HD” and all WT and control samples (20 CAG repeats) as “WT” and trained ML models to identify HD and WT samples ^2,4–6^. Furthermore, from *Langfield et al (2016)* we knew even though there were significant differences between the lower end of the CAG repeat mice with human HD/ JOHD genotypes (80-92 CAG repeats) and the upper end (140-175 CAG repeats), molecular differences were seen in all categories of mice with human HD/ JOHD genotypes, which supports our HD-WT cut-off.

### Machine Learning approach identified miRNAs and mRNAs which can classify WT and HD samples

The goal of this study was to identify potential miRNA and mRNA biomarkers in HD samples, and the initial data exploration helped set realistic queries. From our initial analysis we also found it important to separate the miRNAs and mRNAs, as the DE results showed there was greater variance seen in the mRNAs. The input data consisted of a young miRNAs (519 genes, 40 HD samples, 16 WT samples), aged miRNAs (519 genes, 39 HD samples, 16 WT samples), young mRNAs (532 genes, 40 HD, 16 WT samples) and aged mRNAs (532 gene, 39 HD samples, 16 WT samples).

When finding markers for aged or young samples, a 0.8:0.2 split was used, and when looking for markers which indicate predisposition, the aged samples were trained on, and the young samples were tested on. We initially used RFE, with a 5-fold cross-validation, to identify an optimal minimal number of features, however high numbers of features were identified during aged miRNA training (274), predisposed miRNA training (127), aged mRNA training (140) and young mRNA training (185). RFE selected features were trained using multiple classifiers and the training scores of the miRNAs and mRNAs were usually 1 and testing scores for the miRNAs and mRNAs were respectively between 0.5-0.9 and 0.69-1, indicating a tendency for overfitting (Figure 4A). Instead, we opted to use a robust k-best based feature selection strategy with a smaller number of classifiers (ADA, ET, GNB, RF). These classifiers were selected because they were the best performing classifiers from the RFE based miRNA models. Other classifiers such as Linear SVM and Naïve Bayes also performed well but were not taken forward as they performed poorly for the miRNA-based predisposition question. The number of features found to be responsible for the highest three training scores for each of the four classifiers were taken forward for hyperparameter tuning (Supplementary Table 1). Overall, in contrast to RFE based models, the k-best based models test accuracies were higher and used fewer features for training. Training accuracy, testing accuracy, recall score, precision score, f-score and confusion matrixes were recorded for each model (Figure 4B, Supplementary Table 2).

**Figure 4.**
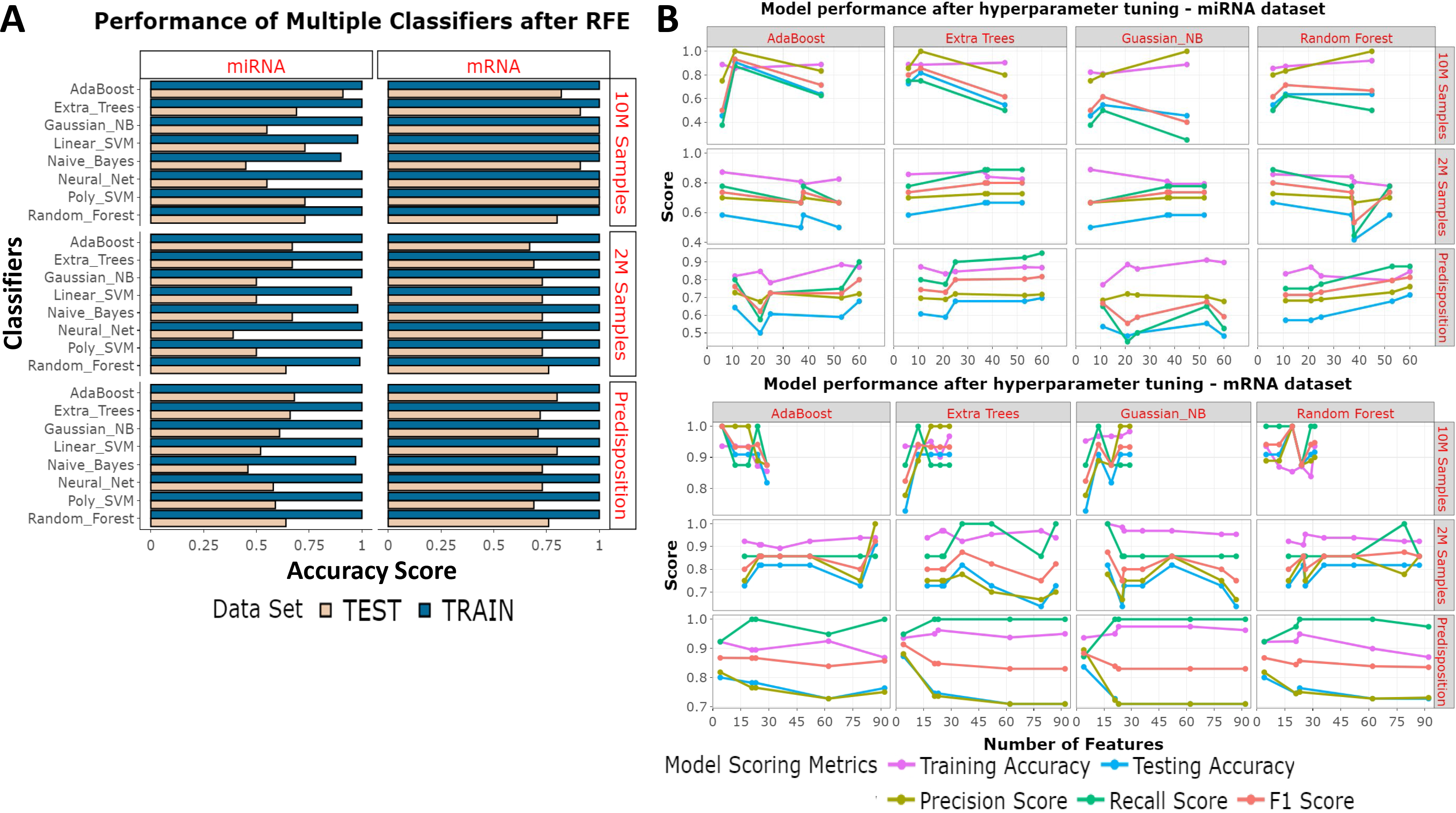
Feature selection approaches. **(A)** RFE was used to identify the optimal features for each question. Eight classifiers were used. The bar chart shows the results from training accuracies (blue) and testing accuracies (beige). (**B)** From a robust k-best based feature identification method the best number of features were taken forward for hyperparameter tuning. Each classifier had each selected set of features hyperparameter tuned. The Training accuracies (purple) testing accuracies (blue), precision scores (yellow-green), recall scores (green) and f1 scores (orange) have been measured from each model.

Best performing models for each question were selected on two criteria –testing accuracies, and precision scores (Table 1). Including good precision scores meant models had to be able to discriminate between WT and HD cases. In cases where multiple classifiers led to the same scores, the classifier which used the least number of features was selected. We also avoided using models built with ET due to high variability of scores seen during the training steps. With reference to the six questions we identified, the best features and classifiers were: a) the 11 miRNAs used to train aged miRNA samples with RF which tested at 85%, b) the 6 miRNAs used to train young miRNA samples with RF which tested at 66%, c) the 60 miRNAs used to train predisposed samples with RF which tested at 71%, d) the 5 mRNAs used to train aged mRNA samples with ADA which tested at 100%, e) the 87 mRNAs used to train young mRNA samples with ADA which tested at 90% and f) the 4 mRNAs used to train predisposed samples with GNB which tested at 83%.

**Table 1.**
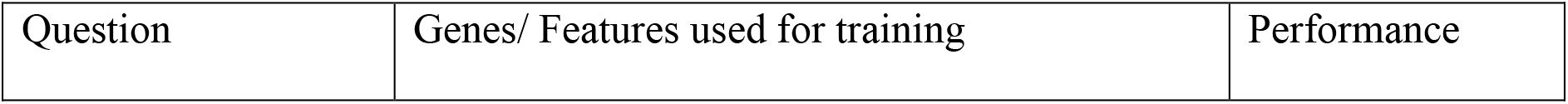
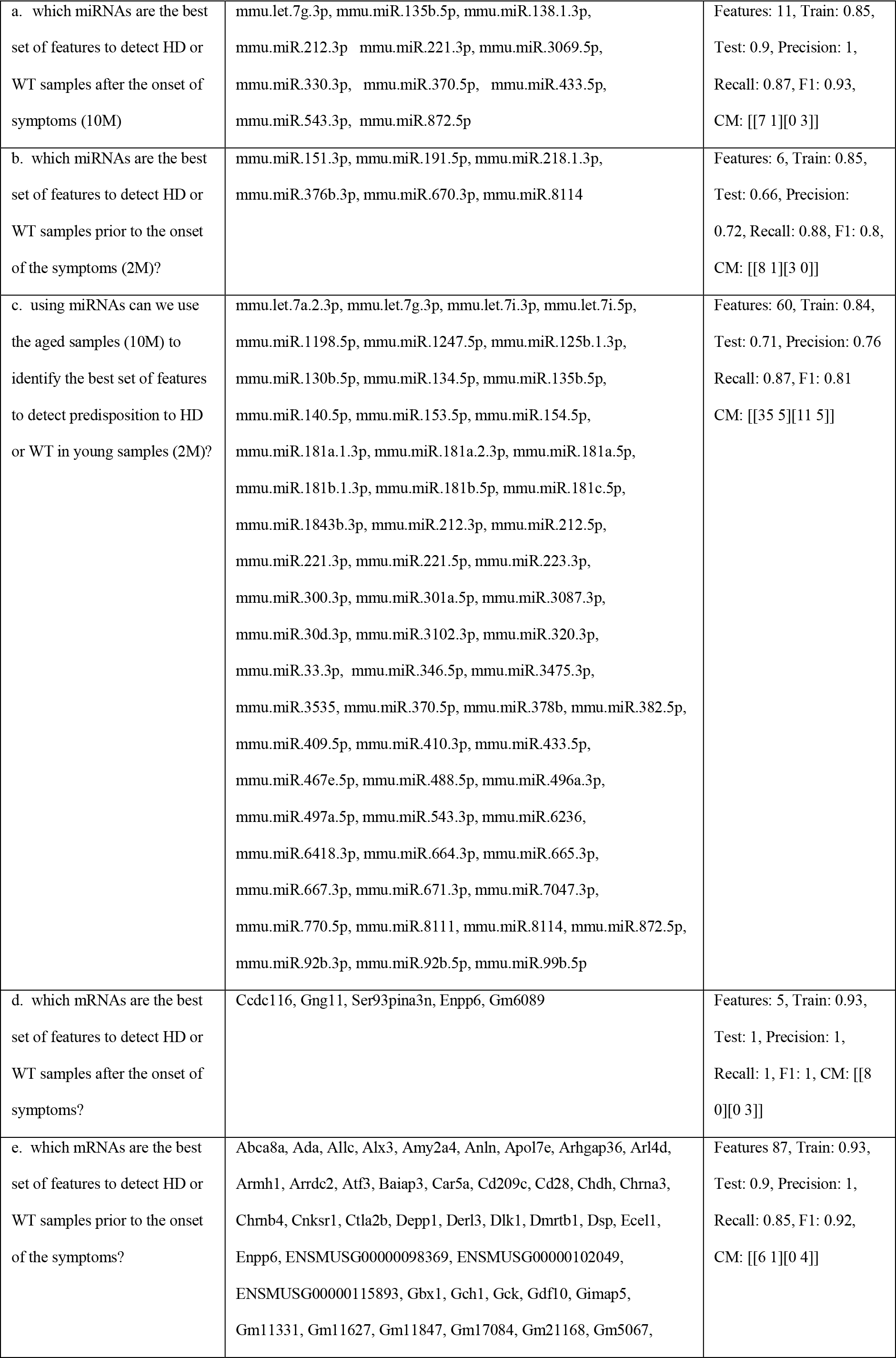
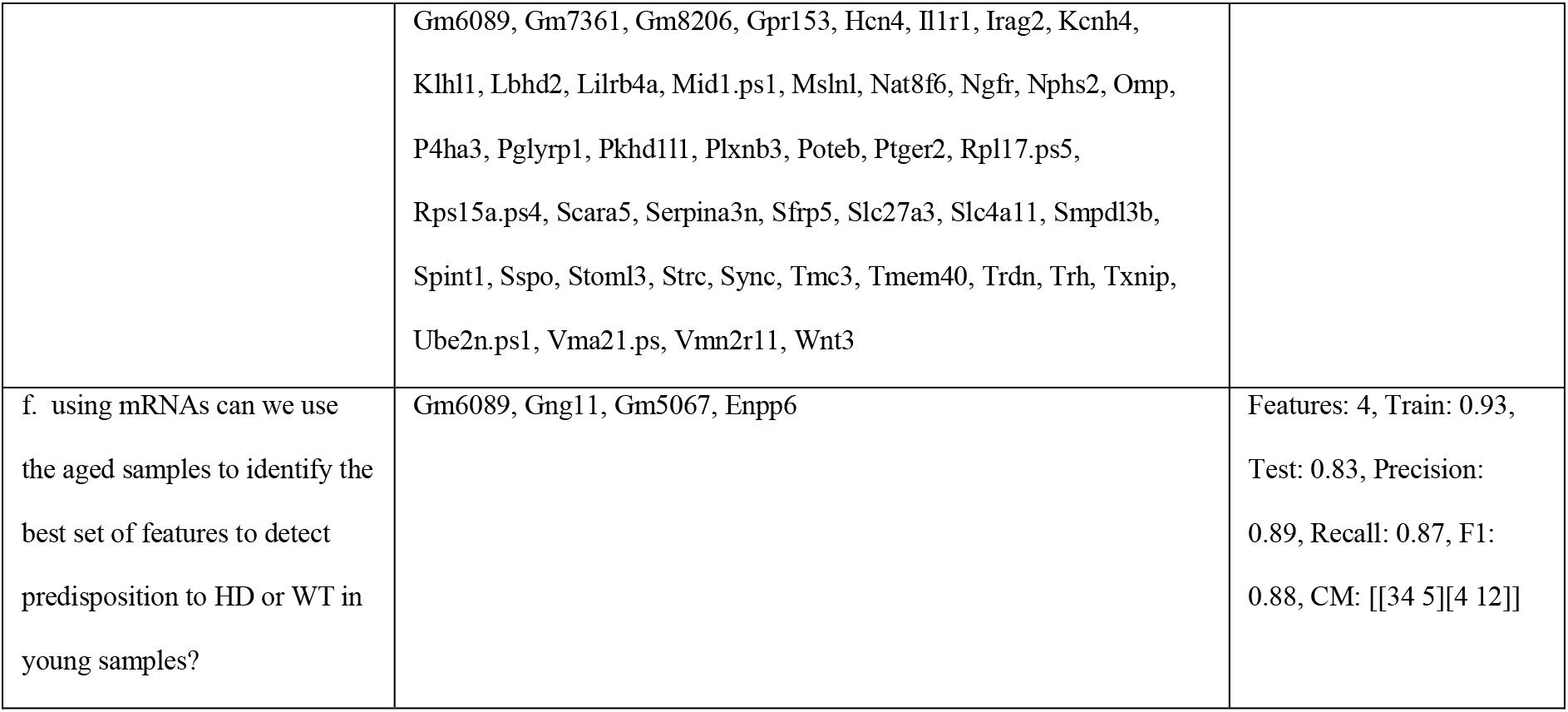
miRNAs/ mRNAs found to be the best features for training models to answer each question. The six questions asked using this dataset are displayed from a-f and the features used for training the best performing models for each question are displayed in alphabetical order. Model scores are also provided, which state the number of features, training accuracies, testing accuracies, precision, recall, f1 and confusion matrices (CM).

### Investigating model performances

AUC plots and PCA plots for the best performing models were created (Figure 5). Unsurprisingly, the mRNA-based models all outperformed their miRNA-based counterparts; most likely because mRNAs showed greater variance than miRNAs which was seen during DE analysis. The miRNA-based 2M was difficult to train with AUC scores of 31%, and the miRNA-based predisposition model achieved an unimpressive, but acceptable AUC score of 63%, though the 10M miRNA-based model had an excellent AUC of 96%. In contrast the mRNA-based models performed well with AUC scores for the mRNA-based 10M, 2M and predisposition models to respectively be 100%, 86% and 92%. The PCA plots for the miRNA-based 2M and predisposition models showed that the model performance was poor when trying to correctly classify true WT samples, meanwhile the PCAs showed the aged miRNA model, and all mRNA-based models were better at classifying WT samples correctly.

**Figure 5.**
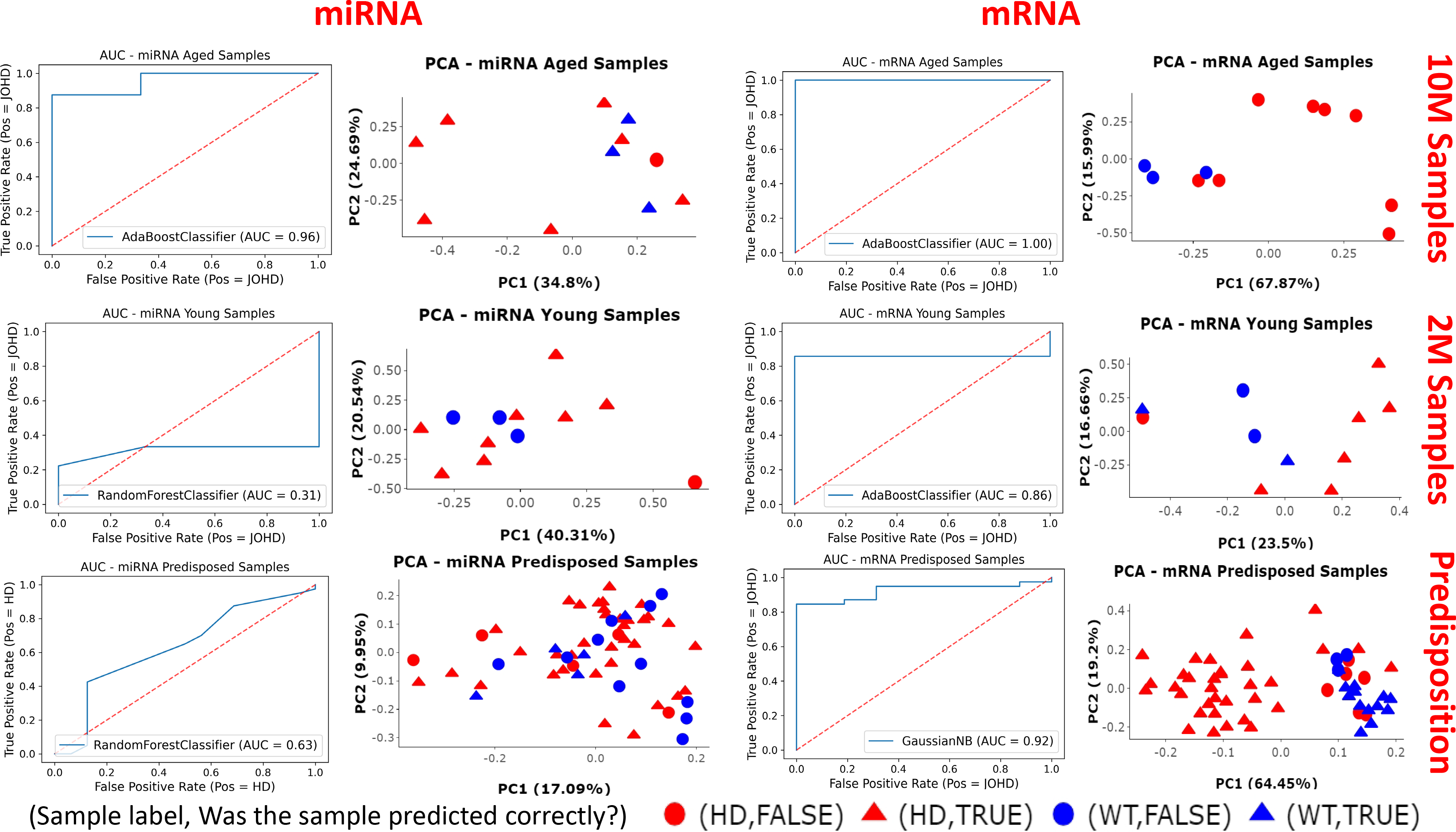
AUC plots and PCA plots of the best performing models. For each of the questions of interest the best performing model was selected for further analysis. The ROC-AUC curves show the performance of the models and PCA plots show if the samples were correctly (triangle) or incorrectly (circle) predicted to be HD (red) or WT (blue).

### Reperforming miRNA-based models using only 140-175 CAG repeat samples

Our miRNA-based models performed worse than their mRNA counterparts, so we decided to re-perform all miRNA models with only the 140 and 175 CAG repeat mouse samples representing the HD cases (Supplementary Table 4). The authors of the dataset stated the higher CAG repeats lead to more severe phenotypes, so a clearer disparity between HD and WT samples may have been achieved if 140-175 CAG repeat samples were used. Unsurprisingly the 10M samples performed well, but generally the 2M and predisposition models performed worse, thus the initial miRNA-based models (WT vs 80-175 CAG repeats) outperformed the reperformed miRNA-based models (WT vs 140-175 CAG repeats).

### Ratification using bio-fluid based transcriptomic datasets for the identified miRNAs

The cortex-based miRNAs used to train our predisposition model needed to be checked if they could be detected while circulating in the bio-fluids to promote them as suitable candidates for use as non-invasive biomarkers for HD research. To investigate this, we downloaded and performed DE on three publicly available miRNA-based HD patient bio-fluid datasets and identified 35 of the 60 miRNAs used to train the miRNA-based predisposition model had known human homologues. Of these 35 miRNAs, six miRNAs were identified to have p.values <0.05 from the blood based datasets. We present the p.values to give a confidence score (lower p.values indicate more confidence of the change in expression observed) and log2fc values to provide a magnitude score (greater the value is away from 0, the greater the expression change between disease and control groups observed). From GSE108395, *hsa-miR-382-5p* had a p.value of 0.0017 and a log2fc value of 0.56, *hsa-miR-770-5p* had a p.value of 0.012 and a log2fc of −0.48, *hsa-miR-154-5p* has a p.value of 0.02 and a log2fc value of 0.43, *hsa-miR-181a-5p* had a p.value of 0.034 and a log2fc value of 0.28 and, *hsa-miR-212-3p* had a p.value of 0.035 and a log2fc value of 0.38. Also, from GSE167630, *hsa-miR-378b* had a p.value of 0.097 and a log2fc value of −0.19.

## Discussion and Conclusion

We analysed a publicly available dataset to train ML models with the aim of identifying potential biomarkers for predisposition to HD. Six questions were asked to achieve this aim (a-f). Questions a and b asked which miRNAs were the best set of features to detect HD or WT genotypes in respectively aged or young samples. Using 11 miRNAs we generated an excellent model for question a which tested at 85% and achieved an AUC score of 96%. In contrast, for question b we used 6 miRNAs to achieve an acceptable testing accuracy of 66% and a poor AUC score at 31%. Questions d and e asked which mRNAs were the best set of features are to detect HD or WT genotypes in respectively aged or young samples. For question d, using 5 mRNAs we created an excellent model which had a testing accuracy and AUC score of 100% and for question e, using 87 mRNAs we created a good model which tested at 90% and had an AUC score of 86%. Questions c and f respectively used miRNAs or mRNAs to ask if we could use aged samples to predict predisposition to HD or WT genotypes in young samples. For question c, with 60 miRNAs we achieved a decent test accuracy of 71% and an acceptable AUC of 63% and for question f, with 4 mRNAs we achieved a good test accuracy of 83% and an excellent AUC of 92%. Overall, for questions a, c, d, e, and f we generated good models with prediction accuracies >70%, and even reached accuracies >90% for questions a, d and e. Generally, we saw poorer performance from miRNA-based models, so perhaps a larger dataset would be required to achieved better results for miRNA-based models. To account for the difference seen between the miRNA-based models we attempted to reperform ML with only 140-175 CAG repeat samples as these had the most severe HD phenotypes ^20,21^. However, these models generally performed far worse, most likely due to the lack of data which left these models poorly trained. Given this we found our assumption of labelling all 80-175 CAG repeat samples as “HD” to be an appropriate decision based on the data available, and this is despite 80-92 CAG repeat mice, 111 CAG repeat mice and 140-175 CAG repeat mice having distinctions in HD development as described by the *Langfelder et al (2016)*. We do believe the models produced in this paper and the sets of gene features found (Table 1) will have use in HD biomarker research. We have recently seen ML becoming a useful resource for investigation of biomarkers for neurodegenerative diseases ^53,54^. Several HD-related studies have been conducted in recent years. For example, a recent ML study identified 66 potential HD genes by combining results from four different statistical/ rules-based algorithms ^55^. Another study used several HD datasets (including the one used in this paper - GSE65776) to identify transcription factors target modules ^56^.

Our predisposition models (questions c and f) were novel and ambitious as the training (10M) and testing (2M) datasets were almost the same size (respectively 55 and 56 samples). However, this was an important question as we know that clinically detectable alterations in the brain and behaviour of HD patients can be seen up to 15 years prior to any the classic HD related motor-loss, indicating early sign of the neurodegenerative disease should be detectable at the gene expression level ^57^. For question f we created an excellent model using four mRNAs. *Enpp6* has been reported as a gene of interest in the hippocampus of HD mice and *Gng11* was associated with 4 of 10 Parkinson’s disease related pathways in a GWAS study, which indicates both genes to be of interest to neurodegenerative research ^58,59^. *Gm5067* and *Gm6089* were predicted protein-coding genes that may have novel roles during HD progression, and both were also identified as features for the best performing model in question e, and the former was identified in question d. In contrast, model we presented for question c achieved an acceptable testing accuracy but an unimpressive AUC score. Even so, as there is substantial interest in miRNAs which can be detected circulating in bio-fluids for use as non-invasive markers to measure the health of neurogenerative disease patients, we further researched the miRNAs used to construct the best performing miRNA-based predisposition model. DE results from three publicly available bio-fluids datasets were mined to identify homologues of the 60 miRNAs selected to train the best performing miRNA-based predisposition model. We found *hsa-miR-154-5p*, *hsa-miR-181a-5p*, *hsa-miR-212-3p, hsa-miR-378b, hsa-miR-382-5p* and hsa-*miR-770-5p* to be DEGs in patient blood plasma samples. There was evidence to link some of these miRNAs with neurodegenerative disorders, such as *miR-181a-5p* and the development of Parkinson’s disease ^60^. Also, *mmu-miR-212-3p* was cited as a potential miRNA of interest in HD mice and hsa-*miR-212* has been linked to neuronal plasticity and cognition ^61,62^. Interestingly, *mmu-miR-378b* was one of the miRNAs highlighted in the data generation study and *hsa-miR-382-5p* has been identified as a miRNA of interest in a Parkinson’s Disease study and an Alzheimer’s disease study, which may indicate it to be a useful miRNA in neurodegenerative disease research ^20,63,64^. No neurodegenerative disease related work was found linked with *hsa-miR-154-5p* or *hsa-miR-770-5p*. Further investigation of these miRNAs could be helpful for HD research and could supplement known HD related miRNAs of interest such as *miR-9* and *miR-9\** which are involved in the regulation of neuron survival antagonist REST ^65^.

It is important to note that age may not have been the only variable of interest - especially in JOHD as large systematic reviews of clinical studies of JOHD patients such as Predict-HD, Enroll-HD and kids-JOHD respectively found age to account for 26%, 59% and 86% of disease development ^17–19^. Thus, age is certainly an important factor in HD/ JOHD, but not the only one. It is also important to ask is how confident can we be that our mouse-based study was suitable for the identification of biomarkers for the benefit of HD patient research. Mouse models have been a popular resource for the study human diseases linked with miRNA expression changes, such as *mmu-miR-140-5p* down-regulation in osteoarthritis or *mmu-miR-21-5p* up-regulation in kidney fibrosis mice ^66,67^. However, mouse-human HD progression does not correlate well, as genotypes which lead to severe neurodegenerative phenotypes via HD related neurotoxicity in humans may resolve as mildly affected/ WT mice, and *Donaldson et al (2021)* reviewed many cases where brain scans and behaviour of mice with human HD/ JOHD genotypes showed no clinical symptoms ^68^. However, it should be noted that many of the poly-glutamine repeat mice in the literature have a mix of CAG and CAA repeats, and CAA repeats have been linked to weaker onset of HD ^69^. The data we used was reported to be from mice that had undergone CAG knock-in experiments ^20,21^. Given this, when creating our ML models, we assumed the 2M mice to not have developed HD yet and the 10M mice to have developed HD symptoms, or to at least have molecular changes associated with HD. Overall, though age was predominantly reported as the major factor in HD research, further research is needed to understand the complexities of HD ^57,70^.

To conclude, we created several good models for mRNA-based samples, including an aged model with 100% accuracy using five mRNAs and an 85% accuracy predisposed model using only four mRNAs - two of which were novel genes. We also highlighted six potential miRNAs which could be markers of predisposition to developing HD. Importantly, these miRNAs were detected in circulating blood samples of HD patients. We believe this analysis study serves as a useful means of hypothesis generation to aid HD researchers and clinicians. The potential of detecting early predisposition biomarkers non-invasively could be of great benefit to neurodegenerative disease research and patient care.

## Author Contributions

KP conceptualised the project, performed all bioinformatics and data preparation. KP, CS generated the machine learning models. KP, DPS contributed equally to writing. KP, CS, DPS approved the submitted version.

## Acknowledgements

We would like to thank a talented UG student Bethany Harley^1^ for her work on gender differences between HD samples. We also thank Professor David Young^1^ for supervisory support and Dr Jamie Soul^1^, Dr Louise Peas^1^ for technical advice during the project.

## Data availability

Preliminary and teaching scripts, plus some exploratory work which was used as a basis for much of the work performed here has been stored in Colleensdan/Predicting-Early-Onset-Huntington-s-Disease. Scripts (DE, *TimiRGeN*, plotting, ML and bio-fluid analysis) and associated data shown in this manuscript are available in Krutik6/Using-Machine-Learning-to-identify-microRNAs-as-biomarkers-for-pre-disposition-to-Juvenile-Onset-Hun.

## References

1. Casella, C., Lipp, I., Rosser, A., Jones, D. K. & Metzler-Baddeley, C. A Critical Review of White Matter Changes in Huntington’s Disease. Movement Disorders vol. 35 Preprint at https://doi.org/10.1002/mds.28109 (2020).

2. Evans, S. J. W. et al. Prevalence of adult Huntington’s disease in the UK based on diagnoses recorded in general practice records. J Neurol Neurosurg Psychiatry 84, (2013).

3. Ohlmeier, C., Saum, K. U., Galetzka, W., Beier, D. & Gothe, H. Epidemiology and health care utilization of patients suffering from Huntington’s disease in Germany: Real world evidence based on German claims data. BMC Neurol 19, (2019).

4. Quarrell, O., O’Donovan, K. L., Bandmann, O. & Strong, M. The prevalence of juvenile Huntington’s disease: A review of the literature and meta-analysis. PLoS Currents Preprint at https://doi.org/10.1371/4f8606b742ef3 (2012).

5. Rosenblatt, A. et al. Age, CAG repeat length, and clinical progression in Huntington’s disease. Movement Disorders 27, (2012).

6. Semaka, A., Collins, J. A. & Hayden, M. R. Unstable familial transmissions of huntington disease alleles with 27-35 CAG repeats (intermediate alleles). American Journal of Medical Genetics, Part B: Neuropsychiatric Genetics 153, (2010).

7. Fusco, F. R. et al. Cellular localization of huntingtin in striatal and cortical neurons in rats: Lack of correlation with neuronal vulnerability in Huntington’s disease. Journal of Neuroscience 19, (1999).

8. Takano, H. & Gusella, J. F. The predominantly HEAT-like motif structure of huntingtin and its association and coincident nuclear entry with dorsal, an NF-kB/Rel/dorsal family transcription factor. BMC Neurosci 3, (2002).

9. Hoffner, G., Kahlem, P. & Djian, P. Perinuclear localization of huntingtin as a consequence of its binding to microtubules through an interaction with β-tubulin: Relevance to Huntington’s disease. J Cell Sci 115, (2002).

10. Caviston, J. P., Ross, J. L., Antony, S. M., Tokito, M. & Holzbaur, E. L. F. Huntingtin facilitates dynein/dynactin-mediated vesicle transport. Proc Natl Acad Sci U S A 104, (2007).

11. Cornett, J. et al. Polyglutamine expansion of huntingtin impairs its nuclear export. Nat Genet 37, (2005).

12. Zuccato, C. et al. Huntingtin interacts with REST/NRSF to modulate the transcription of NRSE-controlled neuronal genes. Nat Genet 35, (2003).

13. McFarland, K. N. et al. MeCP2: A novel huntingtin interactor. Hum Mol Genet 23, (2014).

14. Pan, L. & Feigin, A. Huntington’s Disease: New Frontiers in Therapeutics. Current Neurology and Neuroscience Reports vol. 21 Preprint at https://doi.org/10.1007/s11910-021-01093-3 (2021).

15. Panas, M., Karadima, G., Markianos, M., Kalfakis, N. & Vassilopoulos, D. Phenotypic discordance in a pair of monozygotic twins with Huntington’s disease. Clinical Genetics vol. 74 Preprint at https://doi.org/10.1111/j.1399-0004.2008.01036.x (2008).

16. Georgiou, N. et al. Differential clinical and motor control function in a pair of monozygotic twins with Huntington’s disease. Movement Disorders 14, (1999).

17. Schultz, J. L., Moser, A. D. & Nopoulos, P. C. The association between cag repeat length and age of onset of juvenile-onset huntington’s disease. Brain Sci 10, (2020).

18. Cronin, T., Rosser, A. & Massey, T. Clinical Presentation and Features of Juvenile-Onset Huntington’s Disease: A Systematic Review. Journal of Huntington’s Disease vol. 8 Preprint at https://doi.org/10.3233/JHD-180339 (2019).

19. Squitieri, F., Frati, L., Ciarmiello, A., Lastoria, S. & Quarrell, O. Juvenile Huntington’s disease: Does a dosage-effect pathogenic mechanism differ from the classical adult disease? in Mechanisms of Ageing and Development vol. 127 (2006).

20. Langfelder, P. et al. MicroRNA signatures of endogenous Huntingtin CAG repeat expansion in mice. PLoS One 13, (2018).

21. Langfelder, P. et al. Integrated genomics and proteomics define huntingtin CAG length-dependent networks in mice. Nat Neurosci 19, (2016).

22. Bartel, D. P. MicroRNAs: Target Recognition and Regulatory Functions. Cell vol. 136 Preprint at https://doi.org/10.1016/j.cell.2009.01.002 (2009).

23. Doench, J. G. & Sharp, P. A. Specificity of microRNA target selection in translational repression. Genes Dev 18, (2004).

24. Lee, Y. et al. EMBO J. EMBO Journal vol. 23 Preprint at (2004).

25. Han, J. et al. The Drosha-DGCR8 complex in primary microRNA processing. Genes Dev 18, (2004).

26. Lund, E. & Dahlberg, J. E. Substrate selectivity of exportin 5 and Dicer in the biogenesis of microRNAs. in Cold Spring Harbor Symposia on Quantitative Biology vol. 71 (2006).

27. Tsutsumi, A., Kawamata, T., Izumi, N., Seitz, H. & Tomari, Y. Recognition of the pre-miRNA structure by Drosophila-Dicer-1. Nat Struct Mol Biol 18, (2010).

28. Chendrimada, T. P. et al. TRBP recruits the Dicer complex to Ago2 for microRNA processing and gene silencing. Nature 436, (2005).

29. Rehwinkel, J., Behm-Ansmant, I., Gatfield, D. & Izaurralde, E. A crucial role for GW182 and the DCP1:DCP2 decapping complex in miRNA-mediated gene silencing. RNA 11, (2005).

30. Jonas, S. & Izaurralde, E. The role of disordered protein regions in the assembly of decapping complexes and RNP granules. Genes and Development vol. 27 Preprint at https://doi.org/10.1101/gad.227843.113 (2013).

31. Zekri, L., Huntzinger, E., Heimstädt, S. & Izaurralde, E. The Silencing Domain of GW182 Interacts with PABPC1 To Promote Translational Repression and Degradation of MicroRNA Targets and Is Required for Target Release. Mol Cell Biol 29, (2009).

32. Yamashita, A. et al. Concerted action of poly(A) nucleases and decapping enzyme in mammalian mRNA turnover. Nat Struct Mol Biol 12, (2005).

33. Friedman, R. C., Farh, K. K. H., Burge, C. B. & Bartel, D. P. Most mammalian mRNAs are conserved targets of microRNAs. Genome Res 19, (2009).

34. Valadi, H. et al. Exosome-mediated transfer of mRNAs and microRNAs is a novel mechanism of genetic exchange between cells. Nat Cell Biol 9, (2007).

35. Kumar, S., Vijayan, M., Bhatti, J. S. & Reddy, P. H. MicroRNAs as Peripheral Biomarkers in Aging and Age-Related Diseases. in Progress in Molecular Biology and Translational Science vol. 146 (2017).

36. Rosas, H. D. et al. Regional and progressive thinning of the cortical ribbon in Huntington’s disease. Neurology 58, (2002).

37. Patel, K. et al. TimiRGeN : R/Bioconductor package for time series microRNA– mRNA integration and analysis. Bioinformatics (2021) doi:10.1093/bioinformatics/btab377.

38. Leinonen, R., Sugawara, H. & Shumway, M. The sequence read archive. Nucleic Acids Res 39, (2011).

39. Andrews, S. & others. FastQC: a quality control tool for high throughput sequence data. 2010. Https://Www.Bioinformatics.Babraham.Ac.Uk/Projects/Fastqc/ Preprint at (2010).

40. Friedländer, M. R., MacKowiak, S. D., Li, N., Chen, W. & Rajewsky, N. MiRDeep2 accurately identifies known and hundreds of novel microRNA genes in seven animal clades. Nucleic Acids Res 40, (2012).

41. Langmead, B. Aligning short sequencing reads with Bowtie. Curr Protoc Bioinformatics (2010) doi:10.1002/0471250953.bi1107s32.

42. Patro, R., Duggal, G., Love, M. I., Irizarry, R. A. & Kingsford, C. Salmon provides fast and bias-aware quantification of transcript expression. Nat Methods 14, (2017).

43. Soneson, C., Love, M. I. & Robinson, M. D. Differential analyses for RNA-seq: transcript-level estimates improve gene-level inferences. F1000Res 4, (2015).

44. Love, M. I., Huber, W. & Anders, S. Moderated estimation of fold change and dispersion for RNA-seq data with DESeq2. Genome Biol 15, (2014).

45. Leek, J. T., Johnson, W. E., Parker, H. S., Jaffe, A. E. & Storey, J. D. The SVA package for removing batch effects and other unwanted variation in high-throughput experiments. Bioinformatics 28, (2012).

46. Pedregosa, F. et al. Scikit-learn: Machine learning in Python. Journal of Machine Learning Research 12, (2011).

47. Lemaître, G., Nogueira, F. & Aridas, C. K. Imbalanced-learn: A python toolbox to tackle the curse of imbalanced datasets in machine learning. Journal of Machine Learning Research 18, (2017).

48. Ferraldeschi, M. et al. Circulating hsa-miR-323b-3p in Huntington’s Disease: A Pilot Study. Front Neurol 12, (2021).

49. Dong, X. & Scherzer Clemens. Differential expression analysis of miRNAs expressed in blood and CSF of Huntington’s disease. (Unpublished Processed Data). Gene Expression Omnibus. Retrieved 07/05/2022, from https://www.ncbi.nlm.nih.gov/geo/query/acc.cgi?acc=GSE108398. . (2019).

50. Ritchie, M. E. et al. Limma powers differential expression analyses for RNA-sequencing and microarray studies. Nucleic Acids Res 43, (2015).

51. Kozomara, A., Birgaoanu, M. & Griffiths-Jones, S. MiRBase: From microRNA sequences to function. Nucleic Acids Res 47, (2019).

52. McGeary, S. E. et al. The biochemical basis of microRNA targeting efficacy. Science (1979) 366, (2019).

53. Vamathevan, J. et al. Applications of machine learning in drug discovery and development. Nature Reviews Drug Discovery vol. 18 Preprint at https://doi.org/10.1038/s41573-019-0024-5 (2019).

54. Myszczynska, M. A. et al. Applications of machine learning to diagnosis and treatment of neurodegenerative diseases. Nature Reviews Neurology vol. 16 Preprint at https://doi.org/10.1038/s41582-020-0377-8 (2020).

55. Cheng, J., Liu, H. P., Lin, W. Y. & Tsai, F. J. Identification of contributing genes of Huntington’s disease by machine learning. BMC Med Genomics 13, (2020).

56. Ament, S. A. et al. Transcriptional regulatory networks underlying gene expression changes in Huntington’s disease. Mol Syst Biol 14, (2018).

57. Djoussé, L. et al. Interaction of normal and expanded CAG repeat sizes influences age at onset of Huntington disease. Am J Med Genet 119 A, (2003).

58. Skotte, N. H. et al. Integrative Characterization of the R6/2 Mouse Model of Huntington’s Disease Reveals Dysfunctional Astrocyte Metabolism. Cell Rep 23, (2018).

59. Zhang, M. et al. Genome-wide pathway-based association analysis identifies risk pathways associated with Parkinson’s disease. Neuroscience 340, (2017).

60. Wang, H., Wang, X., Zhang, Y. & Zhao, J. LncRNA SNHG1 promotes neuronal injury in Parkinson’s disease cell model by miR-181a-5p/CXCL12 axis. J Mol Histol 52, (2021).

61. Fukuoka, M. et al. Supplemental Treatment for Huntington’s Disease with miR-132 that Is Deficient in Huntington’s Disease Brain. Mol Ther Nucleic Acids 11, (2018).

62. The miR-132/212 locus: a complex regulator of neuronal plasticity, gene expression and cognition. RNA & DISEASE (2016) doi:10.14800/rd.1375.

63. Nair, V. D. & Ge, Y. Alterations of miRNAs reveal a dysregulated molecular regulatory network in Parkinson’s disease striatum. Neurosci Lett 629, (2016).

64. Lau, P. et al. Alteration of the microRNA network during the progression of Alzheimer’s disease. EMBO Mol Med 5, (2013).

65. Packer, A. N., Xing, Y., Harper, S. Q., Jones, L. & Davidson, B. L. The bifunctional microRNA miR-9/miR-9* regulates REST and CoREST and is downregulated in Huntington’s disease. Journal of Neuroscience 28, (2008).

66. Chau, B. N. et al. MicroRNA-21 promotes fibrosis of the kidney by silencing metabolic pathways. Sci Transl Med 4, (2012).

67. Barter, M. J. et al. Genome-wide microRNA and gene analysis of mesenchymal stem cell chondrogenesis identifies an essential role and multiple targets for miR-140-5p. Stem Cells 33, (2015).

68. Donaldson, J., Powell, S., Rickards, N., Holmans, P. & Jones, L. What is the Pathogenic CAG Expansion Length in Huntington’s Disease. Journal of Huntington’s Disease vol. 10 Preprint at https://doi.org/10.3233/JHD-200445 (2021).

69. Lee, J. M. et al. CAG Repeat Not Polyglutamine Length Determines Timing of Huntington’s Disease Onset. Cell 178, (2019).

70. Gusella, J. F. & MacDonald, M. E. Molecular genetics: Unmasking polyglutamine triggers in neurodegenerative disease. Nat Rev Neurosci 1, (2000).

