## Supplementary material for "Using Machine Learning to identify microRNA biomarkers for predisposition to Huntington’s Disease"

### Supplementary Materials

**Supplementary Figure 1 |** PCAs showing the variance in the HD dataset. Outliers were removed prior to plotting. miRNAs and mRNAs are plotted separately. Top plots show all the data, and bottom plots show either 2m or 10m data points.

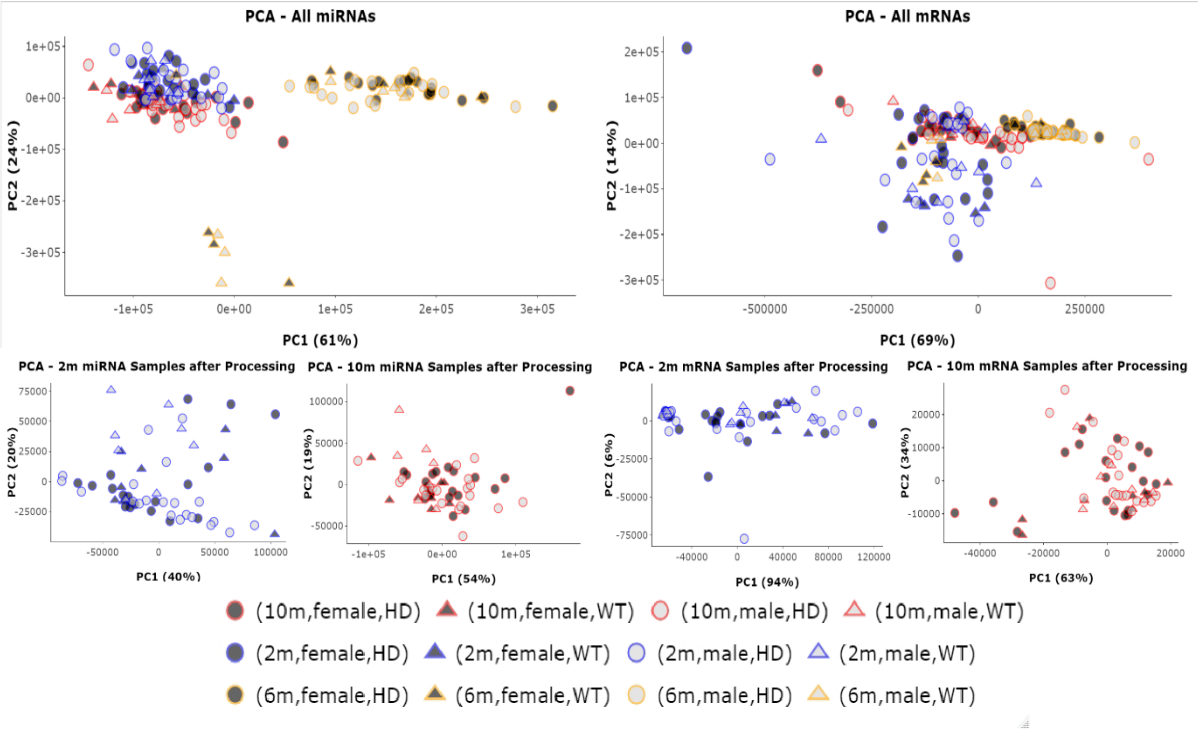

**Supplementary Table 1 |** Best three models from each of the six question. Training accuracies and number of features shown for the best three k-best models for each of the four classifiers. Accuracies shown to two decimal places. Features are ranked from lowest to highest.

| | | | | | | | | | |
|---|---|---|---|---|---|---|---|---|---|
| | Rank | Adaboost | | Gaussian_NB | | Extra_Trees | | RandomForest | |
| miRNA | | training | feat | training | feat | training | feat | training | Feat |
| 10m (aged) | 1st | 0.90 | 11 | 0.88 | 45 | 0.96 | 71 | 0.96 | 45 |
| | 2nd | 0.90 | 11 | 0.88 | 45 | 0.95 | 11 | 0.96 | 45 |
| | 3rd | 0.90 | 11 | 0.88 | 45 | 0.95 | 6 | 0.96 | 45 |
| 2m (young) | 1st | 0.85 | 52 | 0.88 | 6 | 0.93 | 100 | 0.92 | 6 |
| | 2nd | 0.85 | 52 | 0.88 | 6 | 0.93 | 37 | 0.92 | 6 |
| | 3rd | 0.85 | 52 | 0.88 | 6 | 0.92 | 38 | 0.91 | 6 |
| Predisposition | 1st | 0.84 | 11 | 0.92 | 53 | 0.94 | 21 | 0.93 | 53 |
| | 2nd | 0.84 | 11 | 0.92 | 53 | 0.94 | 25 | 0.93 | 53 |
| | 3rd | 0.84 | 11 | 0.92 | 53 | 0.94 | 60 | 0.92 | 53 |
| mRNA | | training | feat | training | feat | training | feat | training | Feat |
| 10m (aged) | 1st | 0.93 | 5 | 0.98 | 29 | 1 | 12 | 1 | 29 |
| | 2nd | 0.93 | 5 | 0.98 | 29 | 1 | 19 | 1 | 29 |
| | 3rd | 0.93 | 5 | 0.98 | 29 | 1 | 24 | 1 | 29 |
| 2m (young) | 1st | 0.95 | 52 | 1 | 17 | 1 | 25 | 1 | 25 |
| | 2nd | 0.95 | 79 | 1 | 17 | 1 | 26 | 1 | 26 |
| | 3rd | 0.95 | 87 | 1 | 17 | 1 | 36 | 1 | 36 |
| Predisposition | 1st | 0.93 | 62 | 0.98 | 92 | 1 | 21 | 0.98 | 92 |
| | 2nd | 0.93 | 62 | 0.98 | 92 | 1 | 23 | 0.98 | 92 |
| | 3rd | 0.93 | 62 | 0.98 | 92 | 0.98 | 23 | 0.98 | 92 |

**Supplementary Table 2 |** K-best based model performances. Results from hyperparameter testing AdaBoost (ADA), Gaussian Naïve Bayes (GNG), Extra Trees (ET) and Random Forest (RF) for each of the six questions asked with this HD dataset. For each model the hyperparameters, training accuracy, testing accuracy, precision score, recall score, F1 score, and confusion matrix (CM) have been displayed. CMs are presented with true positives in the top left quadrant, false positives in the bottom left quadrant, false negatives in the top right quadrant and true negatives in the bottom right quadrant ([[TP FN][FP TN]]). The numbers of features which were selected by the robust k-best method described in the paper, are displayed underneath the classifier name. Each question (best predictors for aged mRNA samples, young mRNA samples, predisposition mRNA samples, aged miRNA samples, young miRNA samples, predisposition miRNA samples) is shaded in grey, each classifier is shaded in light grey and model performances have no shadings. Blue coloured entries were the best performing models.

| miRNA – 10m | | | | | | | |
|---|---|---|---|---|---|---|---|
| AdaBoost | params | training | testing | precision | recall | F1 | CM |
| 6 | lr=0.1, ne=100 | 0.88 | 0.45 | 0.75 | 0.37 | 0.5 | [[3 5] [1 2]] |
| 11 | lr=1, ne=50 | 0.85 | 0.90 | 1 | 0.87 | 0.93 | [[7 1] [0 3]] |
| 45 | lr=0.1, ne=100 | 0.88 | 0.63 | 0.83 | 0.62 | 0.71 | [[5 3] [1 2]] |
| 71 | lr=1, ne=500 | 0.80 | 0.72 | 1 | 0.62 | 0.76 | [[5 3] [0 3]] |
| GaussianNB | params | training | testing | precision | recall | f1 | CM |
| 6 | no smoother | 0.82 | 0.45 | 0.75 | 0.37 | 0.5 | [[3 5] [1 2]] |
| 11 | smoother (0.811130 8307896871) | 0.80 | 0.54 | 0.8 | 0.5 | 0.61 | [[4 4] [1 2]] |
| 45 | no smoother | 0.88 | 0.45 | 1 | 0.25 | 0.4 | [[2 6] [0 3]] |
| 71 | smoother (0.65793 3224657568) | 0.87 | 0.54 | 1 | 0.37 | 0.54 | [[3 5] [0 3]] |
| Extra Trees | params | training | testing | precision | recall | f1 | CM |
| 6 | ne=1000, ms=2 | 0.88 | 0.72 | 0.85 | 0.75 | 0.8 | [[6 2] [1 2]] |
| 11 | ne=10, ms=4 | 0.88 | 0.81 | 1 | 0.75 | 0.85 | [[6 2] [0 3]] |
| 45 | ne=500, ms=2 | 090 | 0.54 | 0.8 | 0.5 | 0.61 | [[4 4] [1 2]] |
| 71 | ne=1000, ms=10 | 0.83 | 0.63 | 1 | 0.5 | 0.66 | [[4 4] [0 3]] |
| RF | params | training | testing | precision | recall | f1 | CM |
| 6 | 10 | 0.85 | 0.54 | 0.8 | 0.5 | 0.61 | [[4 4] [1 2]] |
| 11 | 10 | 0.87 | 0.63 | 0.83 | 0.62 | 0.71 | [[5 3] [1 2]] |
| 45 | 500 | 0.92 | 0.63 | 1 | 0.5 | 0.66 | [[4 4] [0 3]] |
| 71 | 50 | 0.88 | 0.72 | 1 | 0.62 | 0.76 | [[5 3] [0 3]] |
| miRNA – 2m | | | | | | | |
| AdaBoost | params | training | testing | precision | recall | f1 | CM |
| 6 | lr=0.1, ne=100 | 0.87 | 0.58 | 0.7 | 0.77 | 0.73 | [[7 2] [3 0]] |
| 37 | lr=1, ne=1000 | 0.80 | 0.5 | 0.66 | 0.66 | 0.66 | [[6 3] [3 0]] |
| 38 | lr=1, ne=500 | 0.79 | 0.58 | 0.7 | 0.77 | 0.73 | [[7 2] [3 0]] |
| 52 | lr=1, ne=500 | 0.82 | 0.5 | 0.66 | 0.66 | 0.66 | [[6 3] [3 0]] |
| 100 | lr=0.01, ne=500 | 0.79 | 0.75 | 0.75 | 1 | 0.85 | [[9 0] [3 0]] |
| GaussianNB | params | training | testing | precision | recall | f1 | CM |
| 6 | no smoother | 0.88 | 0.5 | 0.66 | 0.66 | 0.66 | [[6 3] [0 3]] |
| 37 | no smoother | 0.81 | 0.58 | 0.7 | 0.77 | 0.73 | [[7 2] |

| | | | | | | | |
|---|---|---|---|---|---|---|---|
| | | | | | | | [3 0]] |
| 38 | no smoother | 0.79 | 0.58 | 0.7 | 0.77 | 0.73 | [[7 2]<br>[3 0]] |
| 52 | no smoother | 0.79 | 0.58 | 0.7 | 0.77 | 0.73 | [[7 2]<br>[3 0]] |
| 100 | no smoother | 0.82 | 0.58 | 0.7 | 0.77 | 0.73 | [[7 2]<br>[3 0]] |
| Extra Trees | params | training | testing | precision | recall | f1 | CM |
| 6 | ne=50, ms=11 | 0.85 | 0.58 | 0.7 | 0.77 | 0.73 | [[7 2]<br>[3 0]] |
| 37 | ne=50, ms=3 | 0.87 | 0.66 | 0.72 | 0.88 | 0.8 | [[8 1]<br>[3 0]] |
| 38 | ne=100, ms=2 | 0.84 | 0.66 | 0.72 | 0.88 | 0.8 | [[8 1]<br>[3 0]] |
| 52 | ne=10, ms=4 | 0.82 | 0.66 | 0.72 | 0.88 | 0.8 | [[8 1]<br>[3 0]] |
| 100 | ne=100, ms=5 | 0.79 | 0.66 | 0.72 | 0.88 | 0.8 | [[8 1]<br>[3 0]] |
| RF | params | training | testing | precision | recall | f1 | CM |
| 6 | 50 | 0.85 | 0.66 | 0.72 | 0.88 | 0.8 | [[8 1]<br>[3 0]] |
| 37 | 50 | 0.84 | 0.58 | 0.7 | 0.77 | 0.73 | [[7 2]<br>[3 0]] |
| 38 | 10 | 0.80 | 0.41 | 0.66 | 0.66 | 0.53 | [[4 5]<br>[2 1]] |
| 52 | 10 | 0.77 | 0.58 | 0.7 | 0.77 | 0.73 | [[7 2]<br>[3 0]] |
| 100 | 10 | 0.80 | 0.58 | 0.7 | 0.77 | 0.73 | [[7 2]<br>[3 0]] |
| miRNA – Predisposition | | | | | | | |
| AdaBoost | params | training | testing | precision | recall | f1 | CM |
| 11 | lr=1, ne=100 | 0.82 | 0.64 | 072 | 0.8 | 0.76 | [[32 8]<br>[12 4]] |
| 21 | lr=1, ne=100 | 0.84 | 0.5 | 0.67 | 0.57 | 0.62 | [[23 17]<br>[11 5]] |
| 25 | lr=1, ne1=00 | 0.78 | 0.60 | 0.72 | 0.72 | 0.72 | [[29 11]<br>[11 5]] |
| 53 | lr=1, ne=1000 | 0.88 | 0.58 | 0.69 | 0.75 | 0.72 | [[30 10]<br>[13 3]] |
| 80 | lr=1, ne=100 | 0.87 | 0.67 | 0.72 | 0.9 | 0.8 | [[36 4]<br>[14 2]] |
| GaussianNB | params | training | testing | precision | recall | f1 | CM |
| 11 | no smoother | 0.77 | 0.53 | 0.68 | 0.65 | 0.66 | [[26 14]<br>[12 4]] |
| 21 | smoother (0.01232<br>8467394420659) | 0.88 | 0.48 | 0.72 | 0.45 | 055 | [[18 22]<br>[7 9]] |
| 25 | smoother (0.0231<br>0129700083159) | 0.86 | 0.5 | 0.71 | 0.5 | 0.58 | [[20 20]<br>[8 8]] |
| 53 | smoother (0.035111<br>91734215131) | 0.91 | 0.55 | 0.70 | 0.65 | 0.67 | [[26 14]<br>[11 5]] |
| 80 | no smoother | 0.89 | 0.48 | 0.67 | 0.52 | 0.59 | [[21 19]<br>[10 6]] |
| Extra Trees | params | training | testing | precision | recall | f1 | CM |
| 11 | ne=100, ms=5 | 0.87 | 0.60 | 0.69 | 0.8 | 0.74 | [[32 8]<br>[14 2]] |
| 21 | ne=1000, ms=12 | 0.83 | 0.58 | 0.68 | 0.77 | 0.72 | [[31 9]<br>[14 2]] |
| 25 | ne=100, ms=7 | 0.84 | 0.67 | 0.72 | 0.9 | 0.8 | [[36 4]<br>[14 2]] |
| 53 | ne=50, ms=2 | 0.87 | 0.67 | 0.71 | 0.92 | 0.80 | [[37 3]<br>[15 1]] |
| 80 | ne=500, ms=4 | 0.86 | 0.69 | 0.71 | 0.925 | 0.81 | [[38 2]<br>[15 1]] |
| RF | params | training | testing | precision | recall | f1 | CM |
| 11 | 10 | 0.83 | 0.57 | 0.68 | 0.75 | 0.71 | [[30 10]<br>[14 2]] |
| 21 | 10 | 0.87 | 0.57 | 0.68 | 0.75 | 0.71 | [[30 10]<br>[13 3]] |
| 25 | 50 | 0.82 | 0.58 | 0.68 | 0.77 | 0.72 | [[31 9]<br>[14 2]] |
| 53 | 50 | 0.79 | 0.67 | 0.72 | 0.87 | 0.79 | [[35 5]<br>[13 3]] |
| 60 | 10 | 0.84 | 0.71 | 0.75 | 0.85 | 0.78 | [[35 5] |

| | | | | | | | |
|---|---|---|---|---|---|---|---|
| | | | | | | | [11 5]] |
| mRNA – 10m | | | | | | | |
| Adaboost | params | training | testing | precision | recall | f1 | CM |
| 5 | lr=0.1, ne=10 | 0.93 | 1 | 1 | 1 | 1 | [[8 0] [0 3]] |
| 12 | lr=0.1, ne=100 | 0.93 | 0.90 | 1 | 0.87 | 0.93 | [[7 1] [0 3]] |
| 19 | lr=0.1, ne=50 | 0.93 | 0.90 | 1 | 0.87 | 0.93 | [[7 1] [0 3]] |
| 24 | lr=0.01, ne=100 | 0.87 | 0.90 | 0.88 | 1 | 0.94 | [[8 0] [1 2]] |
| 29 | lr=0.01, ne=50 | 0.85 | 0.81 | 0.87 | 0.87 | 0.87 | [[7 1] [1 2]] |
| GaussianNB | params | training | testing | precision | recall | f1 | CM |
| 5 | no smoother | 0.95 | 0.72 | 0.77 | 0.87 | 0.82 | [[7 1] [2 1]] |
| 12 | no smoother | 0.96 | 0.90 | 0.88 | 1 | 0.94 | [[8 0] [1 2]] |
| 19 | no smoother | 0.96 | 0.81 | 0.87 | 0.87 | 0.86 | [[7 1] [1 2]] |
| 24 | no smoother | 0.96 | 0.90 | 1 | 0.87 | 0.93 | [[7 1] [0 3]] |
| 29 | no smoother | 0.98 | 0.90 | 1 | 0.87 | 0.93 | [[7 1] [0 3]] |
| Extra Trees | params | training | testing | precision | recall | f1 | CM |
| 5 | ne=500, ms=6 | 0.93 | 0.72 | 0.77 | 0.87 | 0.82 | [[7 1] [2 1]] |
| 12 | ne=100, ms=13 | 0.93 | 0.90 | 0.88 | 1 | 0.94 | [[8 0] [1 2]] |
| 19 | ne=100, ms=5 | 0.95 | 0.90 | 1 | 0.87 | 0.93 | [[7 1] [0 3]] |
| 24 | ne=1000, ms=14 | 0.90 | 0.90 | 1 | 0.87 | 0.93 | [[7 1] [0 3]] |
| 29 | ne=10, ms=10 | 0.96 | 0.90 | 1 | 0.87 | 0.93 | [[7 1] [0 3]] |
| RF | params | training | testing | precision | recall | f1 | CM |
| 5 | 500 | 0.93 | 0.90 | 0.88 | 1 | 0.94 | [[8 0] [1 2]] |
| 12 | 10 | 0.86 | 0.90 | 0.88 | 1 | 0.94 | [[8 0] [1 2]] |
| 19 | 500 | 0.85 | 1 | 1 | 1 | 1 | [[8 0] [0 3]] |
| 24 | 10 | 0.87 | 0.87 | 0.87 | 0.87 | 0.87 | [[7 1] [1 2]] |
| 29 | 10 | 0.83 | 0.90 | 0.88 | 1 | 0.93 | [[8 0] [1 2]] |
| mRNA – 2m | | | | | | | |
| Adaboost | params | training | testing | precision | recall | f1 | CM |
| 17 | lr=0.01, ne=50 | 0.92 | 0.72 | 0.75 | 0.85 | 0.8 | [[6 1] [2 2]] |
| 25 | lr=1, ne=10 | 0.90 | 0.81 | 0.85 | 0.85 | 0.85 | [[6 1] [1 3]] |
| 26 | lr=1, ne=10 | 0.90 | 0.81 | 0.85 | 0.85 | 0.85 | [[6 1] [1 3]] |
| 36 | lr=1, ne=100 | 0.89 | 0.81 | 0.85 | 0.85 | 0.85 | [[6 1] [1 3]] |
| 52 | lr=1, ne=10 | 0.92 | 0.81 | 0.85 | 0.85 | 0.85 | [[6 1] [1 3]] |
| 79 | lr=1, ne=500 | 0.93 | 0.72 | 0.75 | 0.85 | 0.80 | [[6 1] [2 2]] |
| 87 | lr=1, ne=500 | 0.93 | 0.90 | 1 | 0.85 | 0.92 | [[6 1] [0 4]] |
| GaussianNB | params | training | testing | precision | recall | f1 | CM |
| 17 | no smooth | 1 | 0.81 | 0.77 | 1 | 0.87 | [[7 0] [2 2]] |
| 25 | no smooth | 0.98 | 0.63 | 0.66 | 0.85 | 0.75 | [[6 1] [3 1]] |
| 26 | smoother (0.151991108 | 0.96 | 0.72 | 0.75 | 0.85 | 0.8 | [[6 1] [2 2]] |

| | | | | | | | |
|---|---|---|---|---|---|---|---|
| | 29529336) | | | | | | |
| 36 | smoother (0.187381742 2860384)) | 0.96 | 0.72 | 0.75 | 0.85 | 0.8 | [[6 1] [2 2]] |
| 52 | smoother (0.65793322 4657568) | 0.96 | 0.81 | 0.85 | 0.85 | 0.85 | [[6 1] [1 3]] |
| 79 | smoother (0.65793322 4657568) | 0.95 | 0.72 | 0.75 | 0.85 | 0.8 | [[6 1] [2 2]] |
| 87 | no smooth | 0.95 | 0.63 | 0.66 | 0.85 | 0.75 | [[6 1] [3 1]] |
| Extra Trees | params | training | testing | precision | recall | f1 | CM |
| 17 | ne=100, ms=8 | 0.93 | 0.72 | 0.75 | 0.85 | 0.8 | [[6 1] [2 2]] |
| 25 | ne=100, ms=11 | 0.96 | 0.72 | 0.75 | 0.85 | 0.8 | [[6 1] [2 2]] |
| 26 | ne=100, ms=11 | 0.96 | 0.72 | 0.75 | 0.85 | 0.8 | [[6 1] [2 2]] |
| 36 | ne=100, ms=11 | 0.92 | 0.81 | 0.77 | 1 | 0.87 | [[7 0] [2 2]] |
| 52 | ne=10, ms=2 | 0.95 | 0.72 | 0.7 | 1 | 0.82 | [[7 0] [3 1]] |
| 79 | ne=100, ms=7 | 0.96 | 0.63 | 0.66 | 0.85 | 0.75 | [[6 1] [3 1]] |
| 87 | ne=10, ms=13 | 0.93 | 0.72 | 0.7 | 1 | 0.82 | [[7 0] [3 1]] |
| RF | params | training | testing | precision | recall | f1 | CM |
| 17 | 50 | 0.92 | 0.72 | 0.75 | 0.85 | 0.8 | [[6 1] [2 2]] |
| 25 | 500 | 0.90 | 0.81 | 0.85 | 0.85 | 0.85 | [[6 1] [1 3]] |
| 26 | 10 | 0.95 | 0.72 | 0.75 | 0.85 | 0.8 | [[6 1] [2 2]] |
| 36 | 50 | 0.93 | 0.81 | 0.85 | 0.85 | 0.85 | [[6 1] [1 3]] |
| 52 | 50 | 0.93 | 0.81 | 0.85 | 0.85 | 0.85 | [[6 1] [1 3]] |
| 79 | 50 | 0.92 | 0.81 | 0.77 | 1 | 0.87 | [[7 0] [2 2]] |
| 87 | 100 | 0.92 | 0.81 | 0.85 | 0.85 | 0.85 | [[6 1] [1 3]] |
| mRNA – Predisposition | | | | | | | |
| Adaboost | params | training | testing | precision | recall | f1 | CM |
| 4 | lr=0.1, ne=10 | 0.92 | 0.8 | 0.81 | 0.92 | 0.86 | [[36 3] [8 8]] |
| 21 | lr=0.01, ne=50 | 0.89 | 0.78 | 0.76 | 1 | 0.86 | [[39 0] [12 4]] |
| 23 | lr=0.1, ne=10 | 0.89 | 0.78 | 0.76 | 1 | 0.86 | [[39 0] [12 4]] |
| 62 | lr=1, ne=50 | 092 | 0.72 | 0.72 | 0.94 | 0.83 | [[39 0] [15 1]] |
| 92 | lr=1, ne=100 | 0.86 | 0.76 | 0.75 | 1 | 0.85 | [[39 0] [13 3]] |
| GaussianNB | params | training | testing | precision | recall | f1 | CM |
| 4 | no smoother | 0.93 | 0.83 | 0.89 | 0.87 | 0.88 | [[34 5] [4 12]] |
| 21 | smoother (0.01873817 422860384) | 0.95 | 0.72 | 0.72 | 1 | 0.83 | [[39 0] [15 1]] |
| 23 | no smoother | 0.97 | 0.70 | 0.70 | 1 | 0.82 | [[39 0] [16 0]] |
| 62 | no smoother | 0.97 | 0.70 | 0.70 | 1 | 0.82 | [[39 0] [16 0]] |
| 92 | no smoother | 0.96 | 0.70 | 0.70 | 1 | 0.82 | [[39 0] [16 0]] |
| Extra Trees | params | training | testing | precision | recall | f1 | CM |
| 4 | ne=500, ms=7 | 0.93 | 0.87 | 0.88 | 0.94 | 0.91 | [[37 2] [5 11]] |
| 21 | ne=100, ms=2 | 0.95 | 0.74 | 0.73 | 1 | 0.84 | [[39 0] [14 2]] |
| 23 | ne=100, ms=5 | 096 | 0.74 | 0.73 | 1 | 0.84 | [[39 0] [14 2]] |

| | | | | | | | |
|---|---|---|---|---|---|---|---|
| 62 | ne=1000, ms=7 | 0.93 | 0.70 | 0.70 | 1 | 0.82 | [[39 0] [16 0]] |
| 92 | ne=500, ms=3 | 0.95 | 0.70 | 0.70 | 1 | 0.82 | [[39 0] [16 0]] |
| RF | params | training | testing | precision | recall | f1 | CM |
| 4 | lr=0.1, ne=10 | 0.92 | 0.8 | 0.81 | 0.92 | 0.86 | [[36 3] [8 8]] |
| 21 | 100 | 0.924167 | 0.74 | 0.74 | 0.97 | 0.84 | [[38 1] [13 3]] |
| 23 | 10 | 0.94 | 0.76 | 0.75 | 1 | 0.85 | [[39 0] [13 3]] |
| 62 | 10 | 0.89 | 0.72 | 0.72 | 1 | 0.83 | [[39 0] [15 1]] |
| 92 | 10 | 0.87 | 0.72 | 0.73 | 0.97 | 0.83 | [[38 1] [14 2]] |

**Supplementary Table 3 |** Number of DE genes found in each study. The table shows the number of DE genes found from 48 DE analyses, including when genders were combined and when genders were analysed separately. T0 are the numerators and T1 are the denominators for each analysis and miRNAs and mRNAs are shown separately. The numbers of samples per analysis are given after the phenotypic data. M = month.

| DE samples names and numbers | | Number – miRNA | Number - mRNA |
|---|---|---|---|
| T0 | T1 | | |
| CAG20_2M - 8 | WT_2M - 8 | 0 | 6 |
| CAG80_2M - 8 | WT_2M - 8 | 0 | 31 |
| CAG92_2M - 7 | WT_2M - 8 | 0 | 8 |
| CAG111_2M - 8 | WT_2M - 8 | 0 | 13 |
| CAG140_2M - 8 | WT_2M - 8 | 0 | 10 |
| CAG175_2M - 8 | WT_2M - 8 | 0 | 1 |
| CAG20_6M - 8 | WT_6M – 7 | 267 | 5207 |
| CAG80_6M - 8 | WT_6M – 7 | 263 | 5486 |
| CAG92_6M - 8 | WT_6M – 7 | 258 | 4316 |
| CAG111_6M - 8 | WT_6M – 7 | 254 | 5985 |
| CAG140_6M – 8 | WT_6M – 7 | 267 | 5849 |
| CAG175_6M – 7 | WT_6M - 7 | 258 | 6481 |
| CAG20_10M – 7 | WT_10M – 8 | 0 | 0 |
| CAG80_10M – 8 | WT_10M – 8 | 0 | 12 |
| CAG92_10M – 8 | WT_10M – 8 | 0 | 4 |
| CAG111_10M – 8 | WT_10M – 8 | 0 | 9 |
| CAG140_10M – 8 | WT_10M – 8 | 0 | 154 |
| CAG175_10M – 8 | WT_10M - 8 | 34 | 1955 |
| Male_CAG20_2M – 4 | Male_WT_2M - 4 | 0 | 0 |
| Male_CAG80_2M – 4 | Male_WT_2M - 4 | 0 | 2 |
| Male_CAG92_2M – 3 | Male_WT_2M - 4 | 6 | 0 |
| Male_CAG111_2M – 4 | Male_WT_2M - 4 | 1 | 0 |
| Male_CAG140_2M – 4 | Male_WT_2M - 4 | 0 | 0 |
| Male_CAG175_2M – 4 | Male_WT_2M - 4 | 0 | 0 |
| Female_CAG20_2M – 4 | Female_WT_2M - 4 | 1 | 0 |
| Female_CAG80_2M – 4 | Female_WT_2M – 4 | 0 | 4 |
| Female_CAG92_2M – 4 | Female_WT_2M - 4 | 0 | 5 |
| Female_CAG111_2M - 4 | Female_WT_2M - 4 | 2 | 0 |
| Female_CAG140_2M – 4 | Female_WT_2M - 4 | 0 | 1 |
| Female_CAG175_2M - 4 | Female_WT_2M - 4 | 0 | 2 |
| Male_CAG20_6M – 4 | Male_WT_6M – 3 | 334 | 1638 |
| Male_CAG80_6M – 5 | Male_WT_6M – 3 | 321 | 2002 |
| Male_CAG92_6M – 4 | Male_WT_6M – 3 | 6 | 1747 |
| Male_CAG111_6M – 4 | Male_WT_6M – 3 | 313 | 2119 |
| Male_CAG140_6M – 4 | Male_WT_6M – 3 | 317 | 2799 |
| Male_CAG175_6M – 3 | Male_WT_6M – 3 | 319 | 3056 |
| Female_CAG20_6M – 3 | Female_WT_6M - 4 | 288 | 2486 |
| Female_CAG80_6M – 4 | Female_WT_6M – 4 | 294 | 2642 |
| Female_CAG92_6M – 4 | Female_WT_6M - 4 | 276 | 1331 |
| Female_CAG111_6M – 4 | Female_WT_6M - 4 | 283 | 3493 |
| Female_CAG140_6M – 4 | Female_WT_6M - 4 | 293 | 2877 |
| Female_CAG175_6M – 4 | Female_WT_6M - 4 | 289 | 3687 |
| Male_CAG20_10M – 2 | Male_WT_10M - 4 | 3 | 0 |
| Male_CAG80_10M – 4 | Male_WT_10M - 4 | 0 | 2 |

| | | | |
|---|---|---|---|
| Male_CAG92_10M – 4 | Male_WT_10M – 4 | 0 | 7 |
| Male_CAG111_10M – 4 | Male_WT_10M – 4 | 0 | 3 |
| Male_CAG140_10M – 4 | Male_WT_10M – 4 | 0 | 66 |
| Male_CAG175_10M – 4 | Male_WT_10M – 4 | 17 | 537 |
| Female_CAG20_10M – 5 | Female_WT_10M - 4 | 0 | 0 |
| Female_CAG80_10M – 4 | Female_WT_10M – 4 | 0 | 4 |
| Female_CAG92_10M - 4 | Female_WT_10M - 4 | 0 | 5 |
| Female_CAG111_10M - 4 | Female_WT_10M - 4 | 2 | 1 |
| Female_CAG140_10M – 4 | Female_WT_10M - 4 | 1 | 16 |
| Female_CAG175_10M - 4 | Female_WT_10M - 4 | 17 | 796 |

**Supplementary Table 4 |** Model performances from reperforming analysis with Q140-175 as "HD" samples. All stats and nomenclature are the same as with Supplementary Table 2.

| miRNA – 10m | | | | | | | |
|---|---|---|---|---|---|---|---|
| AdaBoost | params | training | testing | precision | recall | F1 | CM |
| 5 | lr=1, ne=500 | 0.88 | 0.85 | 0.66 | 1 | 0.8 | [[2 0]<br>[1 4]] |
| 6 | lr=1, ne=10 | 0.84 | 0.85 | 0.66 | 1 | 0.8 | [[2 0]<br>[1 4]] |
| 8 | lr=1, ne=10 | 0.76 | 1 | 1 | 1 | 1 | [[2 0]<br>[0 5]] |
| 9 | lr=1, ne=10 | 0.76 | 1 | 1 | 1 | 1 | [[2 0]<br>[0 5]] |
| 26 | lr=1, ne=50 | 0.8 | 1 | 1 | 1 | 1 | [[2 0]<br>[0 5]] |
| GaussianNB | params | training | testing | precision | recall | f1 | CM |
| 5 | no smoother | 0.92 | 1 | 1 | 1 | 1 | [[2 0]<br>[0 5]] |
| 6 | no smoother | 0.96 | 1 | 1 | 1 | 1 | [[2 0]<br>[0 5]] |
| 8 | no smoother | 0.92 | 1 | 1 | 1 | 1 | [[2 0]<br>[0 5]] |
| 9 | no smoother | 0.92 | 1 | 1 | 1 | 1 | [[2 0]<br>[0 5]] |
| 26 | no smoother | 1 | 1 | 1 | 1 | 1 | [[2 0]<br>[0 5]] |
| Extra Trees | params | training | testing | precision | recall | f1 | CM |
| 5 | ne=50, ms=2 | 0.92 | 0.85 | 0.66 | 1 | 0.8 | [[2 0]<br>[1 4]] |
| 6 | ne=100, ms=11 | 0.92 | 1 | 1 | 1 | 1 | [[2 0]<br>[0 5]] |
| 8 | ne=100, ms=7 | 0.96 | 1 | 1 | 1 | 1 | [[2 0]<br>[0 5]] |
| 9 | ne=500, ms=5 | 0.96 | 1 | 1 | 1 | 1 | [[2 0]<br>[0 5]] |
| 26 | Ne=500, ms=3 | 0.92 | 1 | 1 | 1 | 1 | [[2 0]<br>[0 5]] |
| RF | params | training | testing | precision | recall | f1 | CM |
| 5 | 50 | 0.96 | 0.85 | 0.66 | 1 | 0.8 | [[2 0]<br>[1 4]] |
| 6 | 50 | 0.96 | 1 | 1 | 1 | 1 | [[2 0]<br>[0 5]] |
| 8 | 50 | 0.92 | 0.71 | 0.5 | 1 | 0.66 | [[2 0]<br>[2 3]] |
| 9 | 100 | 0.96 | 1 | 1 | 1 | 1 | [[2 0]<br>[0 5]] |
| 26 | 500 | 0.96 | 1 | 1 | 1 | 1 | [[2 0]<br>[0 5]] |
| miRNA – 2m | | | | | | | |
| AdaBoost | params | training | testing | precision | recall | f1 | CM |
| 6 | lr=1, ne=10 | 0.88 | 0.28 | 0 | 0 | 0 | [[0 3]<br>[2 2]] |
| 7 | lr=1, ne=10 | 0.77 | 0.28 | 0 | 0 | 0 | [[0 3]<br>[2 2]] |
| 9 | lr=1, ne=50 | 0.77 | 0.14 | 0 | 0 | 0 | [[0 3]<br>[2 2]] |
| 14 | lr=1, ne=50 | 0.81 | 071 | 0.6 | 1 | 0.75 | [[3 0]<br>[2 2]] |
| GaussianNB | params | training | testing | precision | recall | f1 | CM |

| | | | | | | | |
|---|---|---|---|---|---|---|---|
| 6 | smoother(1) | 0.92 | 0.57 | 0.5 | 0.33 | 0.4 | [[1 2]<br>[1 3]] |
| 7 | smoother(1) | 0.96 | 0.42 | 0.33 | 0.33 | 0.33 | [[1 2]<br>[2 2]] |
| 9 | smoother(1) | 0.96 | 0.57 | 0.5 | 0.66 | 0.57 | [[2 1]<br>[2 2]] |
| 14 | smoother(1) | 0.57 | 0.57 | 0.5 | 0.66 | 0.57 | [[2 1]<br>[2 2]] |
| Extra Trees | params | training | testing | precision | recall | f1 | CM |
| 6 | ne=100, ms=5 | 0.92 | 0.28 | 0 | 0 | 0 | [[0 3]<br>[2 2]] |
| 7 | ne=50, ms=10 | 0.92 | 0.57 | 0.5 | 0.33 | 0.4 | [[1 2]<br>[1 3]] |
| 9 | ne=1000, ms=11 | 0.92 | 0.57 | 0.5 | 0.66 | 0.57 | [[2 1]<br>[2 2]] |
| 14 | ne=1000, ms=10 | 0.92 | 0.57 | 0.5 | 0.66 | 0.57 | [[2 1]<br>[2 2]] |
| RF | params | training | testing | precision | recall | f1 | CM |
| 6 | 50 | 0.88 | 0.57 | 0.5 | 0.33 | 0.4 | [[1 2]<br>[1 3]] |
| 7 | 50 | 0.92 | 0.57 | 0.5 | 0.66 | 0.57 | [[2 1]<br>[2 2]] |
| 9 | 10 | 0.85 | 071 | 0.6 | 1 | 0.75 | [[3 0]<br>[2 2]] |
| 14 | 10 | 0.85 | 0.42 | 0.4 | 0.66 | 0.5 | [[2 1]<br>[3 1]] |
| miRNA – Predisposition | | | | | | | |
| AdaBoost | params | training | testing | precision | recall | f1 | CM |
| 3 | lr=0.1, ne=10 | 0.85 | 0.5 | 0.5 | 0.12 | 0.2 | [[2 14]<br>[2 14]] |
| 13 | lr=1, ne=100 | 0.69 | 0.5 | 0.5 | 0.37 | 0.42 | [[6 10]<br>[6 10]] |
| 17 | lr=1, ne1=00 | 0.79 | 0.65 | 0.72 | 0.5 | 0.59 | [[8 8]<br>[3 13]] |
| 29 | lr=0.01, ne=1000 | 0.85 | 0.4 | 0.33 | 0.18 | 0.24 | [[3 13]<br>[6 10]] |
| 53 | lr=1, ne=10 | 0.72 | 0.56 | 0.58 | 0.24 | 0.5 | [[7 9]<br>[5 11]] |
| GaussianNB | params | training | testing | precision | recall | f1 | CM |
| 3 | no smoother | 0.81 | 0.43 | 0.33 | 0.12 | 0.18 | [[2 14]<br>[4 12]] |
| 13 | smoother (1) | 0.9 | 0.5 | 0.5 | 0.37 | 0.42 | [[6 10]<br>[6 10]] |
| 17 | smoother (1) | 0.93 | 0.53 | 0.53 | 0.6 | 0.63 | [[7 9]<br>[6 10]] |
| 29 | smoother (1) | 0.12 | 0.37 | 0.43 | 0.56 | 0.75 | [[9 7]<br>[6 10]] |
| 53 | smoother<br>(0.08111308307896872) | 0.18 | 0.42 | 0.48 | 0.58 | 0.68 | [[12 4]<br>[7 9]] |
| Extra Trees | params | training | testing | precision | recall | f1 | CM |
| 3 | ne=500, ms=14 | 0.75 | 0.43 | 0.37 | 0.18 | 0.25 | [[3 13]<br>[5 11]] |
| 13 | ne=50, ms=7 | 0.85 | 0.46 | 0.45 | 0.31 | 0.37 | [[5 11]<br>[6 10]] |
| 17 | ne=400, ms=3 | 0.84 | 0.46 | 0.45 | 0.31 | 0.37 | [[5 11]<br>[6 10]] |
| 29 | ne=50, ms=4 | 0.87 | 0.46 | 0.47 | 0.5 | 0.48 | [[8 8]<br>[9 7]] |
| 53 | ne=100, ms=5 | 0.86 | 0.56 | 0.55 | 0.62 | 0.58 | [[10 6]<br>[8 8]] |
| RF | params | training | testing | precision | recall | f1 | CM |
| 3 | 10 | 0.75 | 0.37 | 0.35 | 0.31 | 0.33 | [[5 11]<br>[9 7]] |
| 13 | 500 | 0.88 | 0.5 | 0.5 | 0.43 | 0.46 | [[7 9]<br>[7 9]] |
| 17 | 100 | 0.85 | 0.53 | 0.54 | 0.37 | 0.44 | [[6 10]<br>[5 11]] |
| 29 | 50 | 0.85 | 0.5 | 0.5 | 0.43 | 0.46 | [[7 9]<br>[7 9]] |
| 53 | 50 | 0.81 | 0.5 | 0.5 | 0.62 | 0.55 | [[10 6]<br>[10 6]] |
